# Regulatory changes associated with the head to trunk developmental transition

**DOI:** 10.1101/2022.12.18.520961

**Authors:** Patrícia Duarte, Rion Brattig Correia, Ana Nóvoa, Moisés Mallo

## Abstract

Development of vertebrate embryos is characterized by early formation of the anterior tissues followed by the sequential extension of the axis at their posterior end to build the trunk and tail structures, first by the activity of the primitive streak and then of the tail bud. Embryological, molecular and genetic data demonstrate that head and trunk development are significantly different, indicating that the transition into the trunk formation stage involves changes in regulatory gene networks, that might include the acquisition of cell competence to respond to key regulatory factors. Herein, we explored the regulatory changes involved in this developmental transition by assessing the transcriptome and chromatin accessibility profiles from the posterior epiblast region of mouse embryos at embryonic day (E)7.5 and E8.5. We observed changes in various cell processes, including signaling pathways, ubiquitination, ion dynamics and metabolic processes involving lipids that could contribute to the functional switch in the progenitor region of the embryo. Our data also led to the identification of novel mechanisms controlling the differential Wnt functional requirements during head and trunk development. Moreover, we found substantial changes in chromatin accessibility mostly mapping to intergenic regions, indicating a significant switch in the regulatory elements controlling either head or trunk development. In addition, we tested the functional relevance of potential enhancers of *Wnt5a* and *Nr2f2*, identified in the accessibility assays, that reproduced the expression profiles of the target genes. Deletion of these regions by genome editing had limited effect on the expression of those genes, suggesting the existence of redundant enhancers that guarantee robust expression patterns. Overall, this study provides new insights on the regulatory mechanisms that change during the transition from head to trunk development.

**Author Summary:** Vertebrate main body axis is generated sequentially from head to tail. The developmental processes building head and trunk structures are significantly different, and the transition between these two stages requires substantial changes in functional gene regulatory networks. Herein, we explored such changes through genome wide analyses in developing mouse embryos. We observed significant differences in several signaling pathways and in the basic cell machinery, which may interact promoting a functional switch in the differentiating progenitor cells. We also found substantial changes in the accessibility of regulatory elements controlling either head or trunk formation, which conditioned the binding activity of key developmental transcription factors. Overall, our study gives relevant insights into the mechanisms regulating the head to trunk transition that, if disrupted, can lead to embryonic developmental arrest.

## Introduction

During embryonic development the vertebrate body is generated progressively in a head to tail sequence. Although this is a continuous process it occurs in three distinct steps that produce head, trunk and tail structures [1–3]. Each of these stages is characterized by distinct cell dynamics and the generation of a specific set of tissues. For instance, during head development, the embryo establishes the main body axis, lays down the anlage for future brain structures and engages in the process of gastrulation to generate the germ layers [1, 4]. The latter process requires the induction of the primitive streak at the posterior end of the embryo that organizes the emergence of the embryonic endoderm as well as the mesodermal tissues for the head and heart primordia [4]. Genetic analyses in mice have identified key regulators involved in these processes. Some examples include interactions between *Nodal*, *Bmp4* and *Wnt3* to form the primitive streak [5], *Eomes* for the specification of the endodermal layer and mesoderm delamination [6], and *Gata4* and *Gata6* for heart induction [7]. The switch to trunk development is associated with major changes in the growth dynamics of the embryo. It starts elongating the main body axis at the posterior embryonic end by the progressive addition of new tissue produced by the activity of axial progenitors [2,3,8]. This process is associated with the emergence of the neuro-mesodermal cell (NMC) population, the progenitor cells that build the spinal cord and the axial skeleton [2,8–10]. Additional progenitors in the epiblast also lay down the tissues that will contribute to the formation and vascularization of the organs involved in digestive, excretory and reproductive functions of the animal [11]. Similarly to the cells contributing to most embryonic tissues during head development, the progenitors generating trunk structures are also part of the epiblast, which at this stage occupies the posterior end of the embryo [2,9,12]. Also, the primitive streak keeps being the main organizer of progenitor activity during trunk development [2, 8]. However, the regulatory processes undergo major changes. Inactivation of *Tbxt*, the *Cdx* genes, *Wnt3a*, and the combined *Wnt5a* and *Wnt11* loss of function results in embryo truncation at the head to trunk transition, indicating their essential role for trunk development [13–19]. Other factors, like retinoic acid (RA), known to play essential roles during early stages of brain and heart development [20], are also required for trunk development, as silencing this signaling, most typically through inactivation of *Raldh2*, results in developmental arrest at the head to trunk transition [21]. However, the role of RA in this process might differ from that of the other factors, since axial extension can proceed in the absence of this signaling provided that the transition to trunk development is rescued by an acute exogenous RA administration [22].

These observations indicate that the transition into trunk development is associated with a global change in gene regulatory networks, most particularly in the posterior region of the embryo, that switches from gastrulation movements to axial extension. Importantly, many of the factors that control developmental processes during trunk extension are also expressed at earlier stages of development, despite not being required at those stages according to genetic experiments. This indicates that the head to trunk transition also involves a change in the capacity of cells to respond to regulatory factors when entering the trunk formation stage. From a regulatory perspective, this might involve modification of transcription factor (TF) accessibility to their functional targets in the genome.

In this study, we aimed to understand the mechanisms involved in the switch from head to trunk development. For this, we compared transcriptome and chromatin accessibility profiles from the posterior epiblast region of wild type mouse embryos at embryonic day (E)7.5 and E8.5. We observed significant changes in transcriptomic profiles between these two stages. In addition to the expected changes in factors involved in pluripotency and in the *Hox* gene profiles, we observed modifications in a variety of functional groups, including Wnt signaling pathway, ubiquitination systems and lipid metabolic profiles that might interact together to change functional properties at the progenitor region of the embryo. We also observed major changes in chromatin accessibility profiles mostly involving intergenic regions, thus indicating a major switch in regulatory elements controlling head or trunk development, which were associated with changes in the binding activity of key transcription factors. We also found that the absence of RA activity has very limited impact on the changes in chromatin accessibility. In addition, we performed functional tests on specific enhancers identified in the chromatin analyses, including potential regulators of *Wnt5a* and *Nr2f2.* In transgenic reporter experiments these enhancers showed activity compatible with the regulation of the candidate target genes. However, when removed from the genome by edition procedures they had very limited effect on the expression of those genes, indicating the existence of redundant enhancers that provide robustness to the system.

## Results & Discussion

### Transcriptome profile of the posterior epiblast in the developing embryo

To explore the changes in expression of genes involved in trunk formation, we used RNA-seq to obtain the transcriptome profiles from the posterior epiblast region of wild type mouse embryos at E7.5 and E8.5 (Fig 1A), representing respectively the progenitor-containing region of embryos before and after they engage in trunk formation. Principal component analysis separated the samples by timepoint (S1A Fig), revealing the presence of distinct transcriptomic profiles at these two developmental stages. Differential analysis revealed the presence of 2090 genes significantly downregulated, and 1668 genes upregulated at E8.5 relative to E7.5 (Fig 1B and S1 Table). Manual inspection of the list of differentially expressed genes (DEGs) identified downregulation at E8.5 of pluripotency genes, like *Pou5f1* or *Nanog* [23, 24], and genes involved in the initial establishment of the body axis and germ layers like *Tdgf1* (*Cripto*), *Nodal* or *Eomes* (Fig 1C) [6,25,26]. Conversely, activation of central and posterior *Hox* genes was clearly observed at E8.5 (Fig 1D). These findings fit with expression patterns reported for these genes, thus serving as an initial validation of our approach.

**Figure 1.**
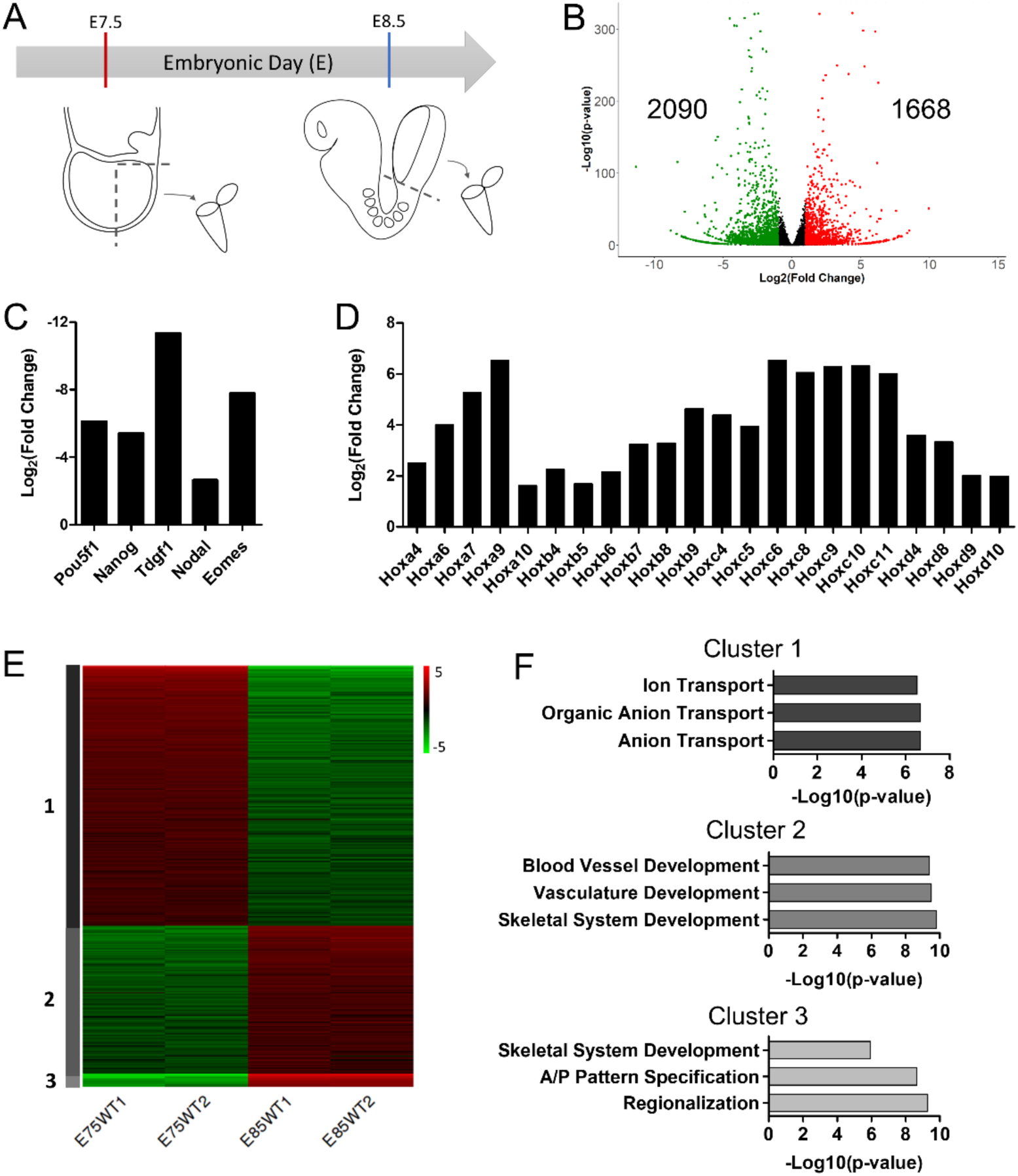
Transcriptomic changes in the posterior epiblast associated with the head to trunk transition. (A) Schematic representation of sample collection from the posterior epiblast region of mouse embryos at E7.5 and E8.5. (B) Volcano plot of RNA-seq gene expression (|Log_2_(Fold Change)|≥1 & p-value < 0.05). Significantly upregulated genes at E8.5 are in red, downregulated at E8.5 are in green and non-significant in black. (C-D) Gene expression of key pluripotent and early developmental genes (C) and Hox genes (D). (E) K-means clustering of the 1000 most variable genes. Cluster 1: 616 genes; Cluster 2: 352 genes; Cluster 3: 32 genes. (F) Top 3 GO terms from biological processes associated with Cluster 1, 2 and 3.

K-means clustering of the top 1000 most variable genes produced three clusters with distinct gene expression dynamics (Fig 1E and 1F). Cluster 1 includes genes that became downregulated at E8.5; genes in this cluster are enriched in gene ontology (GO) terms related to anion and ion transport. Interestingly, a similar decrease in expression of genes enriched for ion transport and homeostasis has been described at the whole embryo level during the same stages analyzed here [27], further suggesting an important role for changes in ion transport profiles during early embryonic development. The full implication of this finding remains elusive. The control of ion fluxes has been implicated in patterning processes [28, 29], including early stages in the establishment of left-right asymmetry associated with node activity [30, 31]. They also have been shown to control cell processes involved in cell migration, cell proliferation and autophagy [32–34]. Focused experimental approaches will be required to explore if the drastic changes in ion transporter profiles observed in the progenitor-containing region during the head to trunk transition play a relevant role in the transition. Cluster 2 comprises genes moderately upregulated at E8.5, mostly associated with skeletal system, vasculature, and blood vessel development. Finally, cluster 3 is composed of genes strongly upregulated at E8.5. Genes in this cluster are enriched in skeletal system development, anterior/posterior pattern specification, and regionalization.

To get a closer image of the changes associated with the transition from head to trunk development, we built a protein-protein interaction (PPI) network (Fig 2A and S1 Network) based on the differentially expressed genes between E7.5 and E8.5, as obtained from StringDB [35]. To focus our analysis on the most relevant interactions, we computed the metric backbone of this PPI network [36], which removed all redundant interactions and has been shown to help identifying genes and interactions responsible for core cellular programs [37]. Next, we identified structurally coherent network modules using LowEnDe [38], with an in-house developed algorithm based on spectral decomposition and information theory. Our interpretation is that these network modules may represent core development functions that are responsible for key aspects of the head to trunk transition.

**Figure 2.**
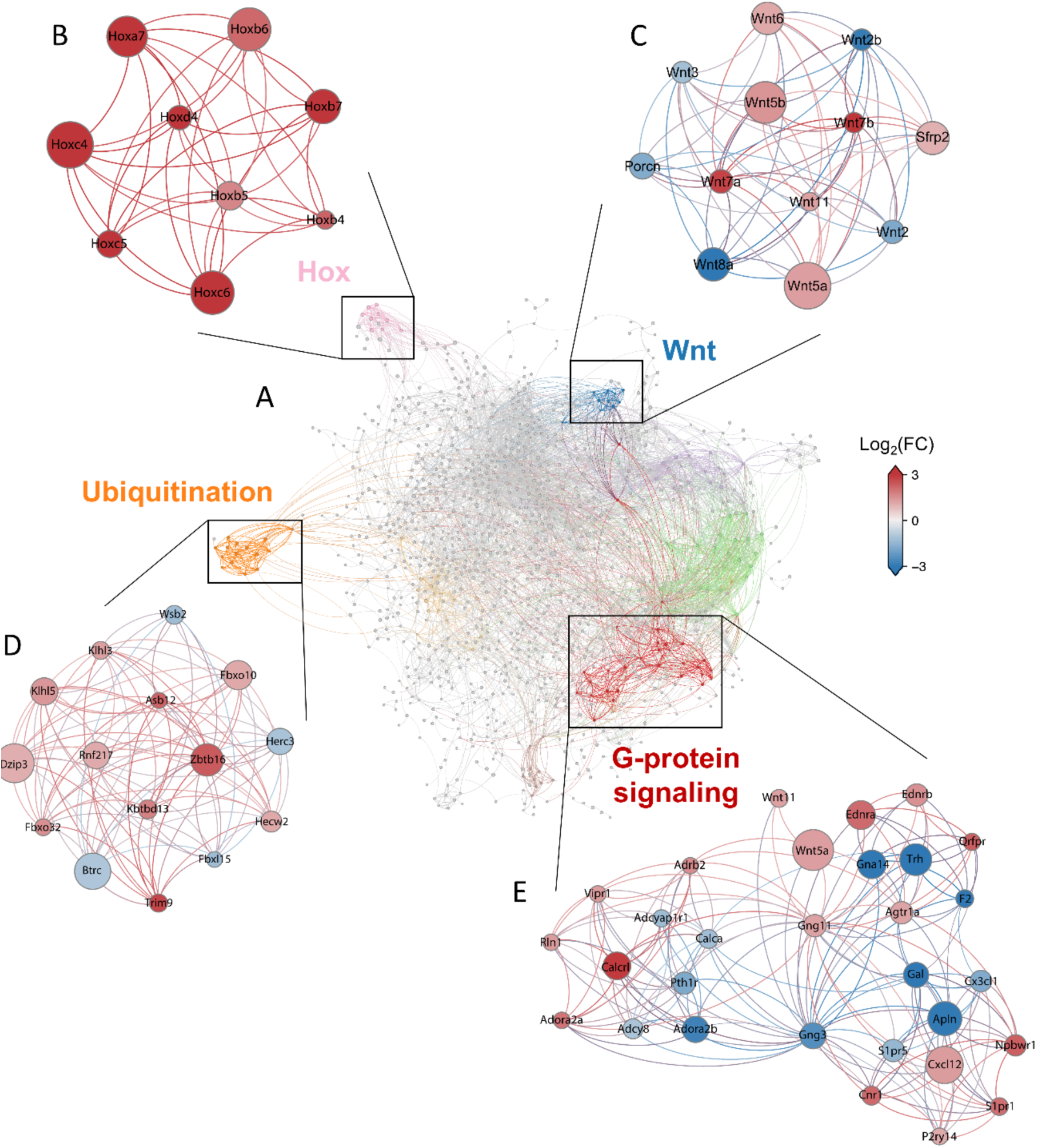
Interaction networks reveal changes in various functional modules. (A) Protein-protein interaction network based on the differentially expressed genes between E7.5 and E8.5. Colored clusters represent structurally coherent network modules identified using LowEnDe [38]. (B-D) Expanded versions of the Hox (B), Wnt (C), G-protein coupled receptor signaling (D) and ubiquitination (E) modules are shown to highlight the genes included in each. Nodes are colored by Log_2_(Fold Change), node size by Log_2_(CPM). Significantly upregulated genes at E8.5 are in red, downregulated genes at E8.5 are in blue.

One of the resulting clusters comprised the *Hox* genes (Fig 2B) that we had already identified in our manual inspection of the differentially regulated genes, thus serving again as an internal validation of the approach. Another prominent cluster was associated with ubiquitination processes (Fig 2D) enriched in genes encoding for E3 ligases, the key determinants of substrate specificity of the ubiquitin proteasome system [39]. This cluster contains a mix of up and downregulated genes, suggesting a switch in global ubiquitination patterns during the head to trunk transition that could impact general cellular functions by changing the availability of components involved in those processes. The large number of canonical Wnt signaling components that have been shown to be regulated by ubiquitination, including β-catenin, Fzd, Lrp6, Dvl, APC and Axin, [40–44], suggests that changes in ubiquitination patterns might lead to functional modifications in Wnt signaling during the head to trunk transition (see also below). Particularly interesting in this module is Btrc, known to promote β-catenin ubiquitination and its subsequent degradation [45–47], which has been shown to also interact with components of several other signaling pathways and regulators of cell proliferation [48–50]. Changes in *Btrc* levels might therefore affect interactions and activity balance between different signaling pathways, eventually impacting their global functional output. Indeed, the PPI network also identified several clusters composed of genes involved in different signaling pathways, indicating the existence of a substantial change in the signaling activities governing cell function when embryos engage in trunk formation.

One of those signaling-related clusters particularly prominent in the PPI network was composed of genes involved in G-protein coupled receptor signaling (Fig 2E). This cluster included both up-and down-regulated genes, suggesting a significant switch in G-protein-dependent signaling during the transition. In addition to changes in ligands and receptors associated with G-protein-mediated signaling, like Ednra, Ednrb, Adora2a, Calcrl or Agtr1a, whose upregulation could be related to the development of the vascular system [51–54], or Cxcl12, which could be related to germ cell migration [55], other changes in the G-protein-associated cluster could indicate a more general functional switch in the basic G-protein machinery. For instance, cluster analysis indicates a switch in the gamma subunits of the heterotrimeric G protein complexes, from Gng3 to Gng11, which could impact the selection of the pathways supported by the complex.

Those general changes in G protein-mediated signaling might play a role in the functional changes associated with Wnt signaling during the head to trunk transition. In particular, the PPI network showed connections between the G-protein cluster and *Wnt5a* and *Wnt11*. This connection might expose a regulatory switch, considering that these Wnt factors are known to signal through non-canonical pathways [56–58] and their activity is essential when the embryo enters trunk development [18, 19]. It will be therefore interesting to determine whether the changes observed in the molecular composition of the G-protein signaling cluster from E7.5 to E8.5, promotes activation of the non-canonical Wnt/Ca^2+^ pathway by Wnt5a and Wnt11 [58] when the embryo engages in axial extension. A more prominent involvement of the non-canonical Wnt signaling downstream of Wnt5a when entering trunk development was also suggested by the upregulated *Sfrp2* expression at E8.5 (Fig 2C), since Sfrp2 redirects Wnt signals from Fz7 to Ror2, stabilizing the Wnt5a-Ror2 complexes that mediate Wnt5a activity during body axis development [59, 60]. The possible involvement of *Sfrp2* in this process is also supported by genetic data showing its requirement during trunk axial extension redundantly with *Sfrp1* [61].

Another of the relevant changes in Wnt signaling associated with the head to trunk transition is the switch from *Wnt*3 to *Wnt3a* functional dependency [17, 47]. In our datasets, *Wnt3* was downregulated at E8.5, fitting with its functional dynamics. *Wnt3a* expression levels, however, did not change from E7.5 to E8.5. This contrasts with the known *Wnt3a* functional requirements, as it is essential during trunk development but seems to be either inactive or functionally limited at earlier developmental stages given its inability to replace for *Wnt3* [47]. This could suggest that stimulation of Wnt3a functional activity during axial extension might result from expression changes in additional factors modulating Wnt signaling at different levels of the pathway. This is an intriguing possibility considering the differential effects that stabilization of Axin2 had on canonical Wnt signaling in different areas of the embryo [62]. In particular, it was shown that in embryos carrying an Axin2 allele (*canp*) coding for a Axin2 protein with increased stability, canonical Wnt signaling was suppressed in the primitive streak during gastrulation and in the neural crest at later stages but was strongly up-regulated specifically in the progenitor zone of E8.5 embryos, eventually negatively impacting axial extension [62]. These observations suggested fundamental changes in Wnt signaling as embryos engage in axial extension, mostly involving the canonical pathway. A prominent candidate to be involved in differential Wnt regulation is *Porcn*, which codes for a molecule that introduces an essential palmitoleoyl moiety into a highly conserved serine residue of the Wnt ligands [63, 64]. Given the essential role of *Porcn* during gastrulation [65], it was somewhat surprising to find a reduction of *Porcn* expression levels at E8.5, which was also observed in the expression patterns reported for this gene [65]. Whether this reduction plays a role in the Wnt signaling switch associated with the head to trunk transition is unclear. Intriguingly, it has been reported that pharmacological inhibition of Porcn impacted differently canonical and non-canonical Wnt signaling in a cell line assay [66], suggesting both that the Porcn-mediated modification might not be a universal requirement for Wnt signaling and that *Porcn* expression levels might control the balance between canonical and non-canonical pathways.

### Wnt signaling dependency on Porcn during axial extension

We tested the effect of blocking Porcn activity on axial extension by incubating E8.5 embryos *in vitro* in the presence or absence of the Porcn inhibitor IWP-01. Our culture conditions allowed normal progression of development, with the embryos attaining typical E9.5 morphology within 24 hours of incubation (Fig 3). The presence of the inhibitor affected development in different ways. The brain structures were seriously reduced in size, likely affecting mainly the midbrain and anterior hindbrain structures, which also led to a substantial reduction in migratory cranial neural crest cells (Fig 3A – Crabp1 panel). These features are consistent with the inhibition of Wnt1 signaling [67], thus serving as an internal control for IWP-01 activity. IWP-01 treated embryos underwent considerable extension at the caudal embryonic end, although they eventually became truncated. Uncx4.1 expression indicated the presence of paraxial mesoderm along the whole anterior posterior axis, presenting fairly normal-looking somites for a considerable extent of the trunk, but losing segmental patterns towards the end of the axis (Fig 3B). The *Uncx4.1* signal almost reached the caudal embryonic end, indicating that the presomitic mesoderm (PSM) was strongly reduced or absent, an idea also suggested by the lack of *Msgn1* signal (Fig 3A). *Sox2* expression indicated that IWP-01-treated embryos also developed a spinal cord, morphologically normal at the axial levels containing identifiable somites and becoming a wider flattened structure in the region containing the disorganized *Uncx4.1* expression (Fig 3B – middle panel). Importantly, even in the area showing abnormal neural and paraxial mesodermal patterns, IWP-01 treated embryos contained a single neural tube. The axial truncation in the context of a disorganized paraxial mesoderm and enlarged spinal cord could indicate an exhaustion of NMCs derived from accelerated progenitor differentiation at the expense of self-renewal. The lack of *Cdx2* expression at the caudal end of IWP-01-treated embryos is consistent with this hypothesis (Fig 3A). Interestingly, *Shh* expression showed that the notochord also became truncated in the region where the paraxial mesoderm and the neural tube lose normal patterns (Fig 3B – right panel). *Tbxt* expression was reduced to a small spot beneath the neural tube (Fig 3A), roughly corresponding to the position of the caudal end of *Shh* expression, indicating that it could represent the posterior end of the notochord.

**Figure 3.**
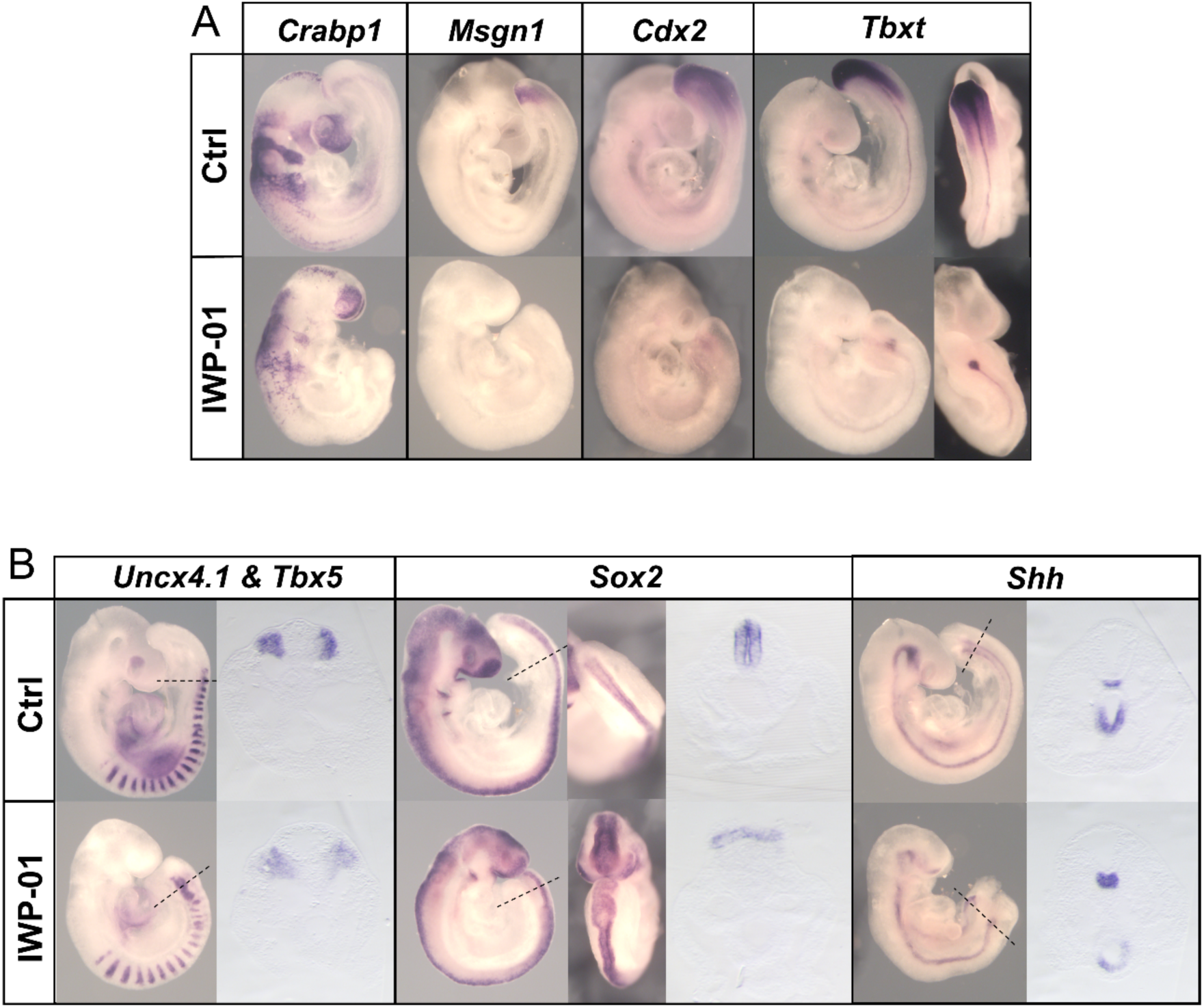
Impact of Porcn activity during axial elongation. Embryos were cultured for 24hrs (E8.5 to E9.5) in the presence or absence of Porcn inhibitor, IWP-01. (A) Whole-mount *in situ* hybridization with *Crabp1, Msgn1, Cdx2* and *Tbxt* probes. (B) Whole-mount *in situ* hybridization with *Uncx4.1* & *Tbx5*, *Sox2* and *Shh* probes, and respective transverse sections.

Even considering that IWP-01 might not block Porcn activity completely, our data indicates that during axial extension Wnt signaling involves a combination of Porcn dependent and independent activities. This contrasts with the essential role of Porcn during gastrulation [65]. Interestingly, the axial level at which the paraxial and spinal cord of IWP-01-treated embryos lose normal patterns roughly corresponds to the level when the embryo starts the transition into tail bud-dependent elongation, thus suggesting different requirements for the control of epiblast-driven and tail bud-dependent axial elongation. The differences between the reported phenotype of *Wnt3a* mutant embryos and the IWP-01 treated embryos indicate that during axial elongation Wnt3a signaling might include Porcn independent activities, an effect previously observed in a cell culture context [68]. The presence of both neural and paraxial mesodermal tissues throughout the AP axis of IWP-01 treated embryos differs from the duplicated neural tubes replacing the paraxial mesoderm characteristic of the *Wnt3a* mutant embryos [69], thus indicating that the Wnt3a activity that modulates NMC cell fate might be Porcn-independent. The malformations observed at the caudal end of the IWP-01-treated embryos suggest that the required equilibrium between differentiation and self-renewal of NMC cells might also entail proper balance of Porcn-dependent and Porcn independent Wnt activities.

### Chromatin accessibility landscape of the posterior epiblast in the developing embryo

To understand the regulation behind the changes observed in gene expression, we mapped global chromatin accessibility profiles. For this, we performed ATAC-seq [70] from tissues of the same regions and timepoints as those used for RNA-seq. Principal component analysis separated the samples by timepoint (S1B Fig), indicating the presence of distinct chromatin accessibility profiles at these two developmental stages. Both E7.5 and E8.5 datasets had a similar chromosomal distribution of accessible regions, with two thirds mapping to promoters and about 20% to intergenic regions (Fig 4A). Differential analysis of the two datasets identified 18197 regions with increased chromatin accessibility (open regions), and 11087 with decreased accessibility (closed regions) at E8.5 relative to E7.5 (Fig 4B and S2 Table). Interestingly, the differentially accessible peaks followed a distribution different to that observed for the individual datasets, with most peaks (57%) mapping to intergenic regions, 14% to introns and the contribution of promoters being reduced to around 21% (Fig 4A). This suggests that the transition between these developmental stages is to a large extent associated with a switch in regulatory elements. In addition, the finding that there are around ten times more genomic regions changing accessibility profiles than differentially expressed genes suggests a high complexity in the regulatory mechanisms controlling the transcriptional switch associated with the head to trunk transition.

**Figure 4.**
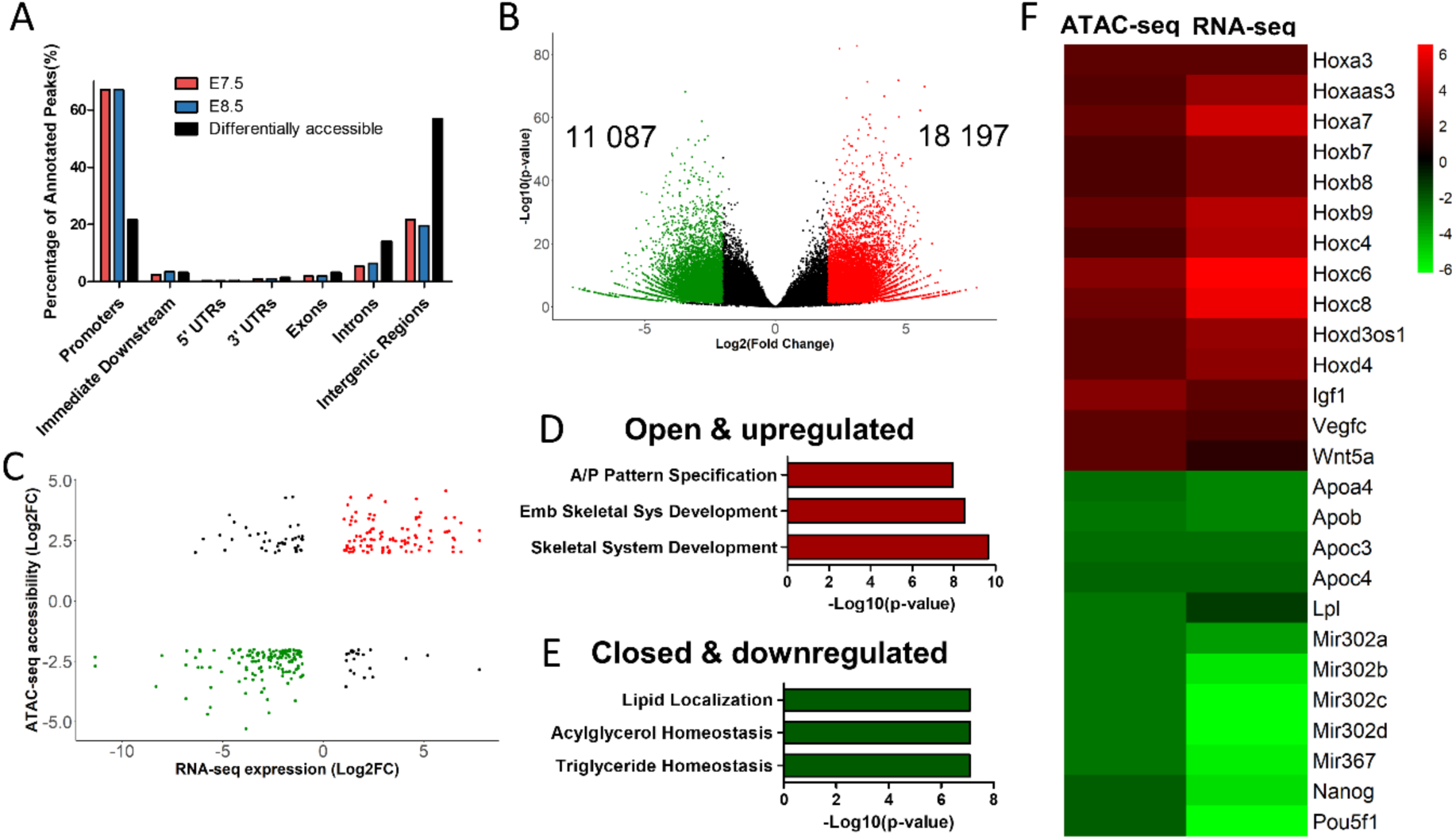
Integration of genome accessibility and gene expression data. (A) Genomic distribution of ATAC-seq peaks identified at E7.5 (red), E8.5 (blue) and distribution of only the differentially accessible peaks (black). (B) Volcano plot of ATAC-seq peaks (|Log_2_(Fold Change)|≥2 & p-value < 0.05). Significantly open regions at E8.5 in red, closed regions at E8.5 in green and non-significant in black. Scatterplot showing correlation between genomic accessibility and gene expression. Significantly accessible and upregulated genes in red, closed and downregulated genes in green. (D) Top 3 GO biological process terms of positively regulated genes at E8.5 (red group in C), Abbreviations: Emb, Embryonic; Sys, System, A/P, Anterior/Posterior. (E) Top 3 GO biological process terms of negatively regulated genes at E8.5 (green group in C) (F) Heatmap of Log_2_(Fold Change) of ATAC-seq and RNA-seq signals.

From the regions showing differential accessibility, only 1418 could be associated with an annotated gene within 5 kb. Integrative analysis of transcriptomic and chromatin dynamics by crossmatching these 1418 regions with the differentially expressed genes (n=3758) identified 300 genes in common, of which 238 showed consistent regulation at both chromatin and transcriptomic levels (Fig 4C) (i.e., upregulated transcripts close to regions that became accessible or downregulated transcripts close to regions that lost accessibility). The remaining 62 regions might represent inhibitory elements. These observations indicate that only a very small proportion of the regions that change accessibility during the head to trunk transition are predicted to control the closest annotated transcriptional unit, thus further complicating the understanding of the regulatory processes controlling the head to trunk transition. Analysis of GO terms of this restricted group revealed an enrichment in anterior/posterior pattern specification and skeletal system development, in genes which are both accessible and upregulated at E8.5 (Fig 4D). These include several *Hox* genes, most particularly those of central and posterior paralog groups (Fig 4F), which might reflect the activation of enhancers within the *Hox* clusters upon sequential global opening of the clusters during axial extension [71]. The group of less accessible and downregulated genes include genes related to stem cell pluripotency and proliferation (Fig 4F), like the already mentioned *Pou5f1* and *Nanog.* This is consistent with the known position of relevant regulatory regions for these genes [72, 73]. This group also included the *miR-302/367* cluster, important for stem cell maintenance and repression of cell differentiation [74].

GO terms of the less accessible and downregulated genes were enriched for triglyceride homeostasis and lipid metabolism (Fig 4E), including several *Apo* genes as well as *Lpl*, that catalyzes the hydrolysis of triglycerides (Fig 4F). These observations indicate that the head to trunk transition is associated with changes in lipid metabolism, which have the potential to impact the activity of various signaling pathways. For instance, lipid modifications have been shown to be essential to generate functionally competent Wnt and Hedgehog molecules [75, 76]. In the case of Wnt ligands, they contain several lipidic modifications, including the above-mentioned palmitoleoylation, which have been shown to affect differently the functional activity of different Wnt molecules [66, 77] and, as already discussed above, could be involved in the implementation of the functional switch in Wnt signaling associated with the head to trunk transition. Interestingly, in Drosophila embryos lipid-modified Hedgehog and Wingless require association with lipoproteins for long-range spreading of their activity [78], and Wnt5a has also been shown to associate with lipoprotein particles for long distance regulation of hindbrain development [79]. In our datasets, several genes encoding for lipoprotein components are downregulated at E8.5. While this could be related to changes in the transport of lipid nutrients to the developing embryo, as shown for Apob during mouse embryogenesis [80], it could also impact Wnt and Hedgehog activities by determining the spatial range of their activity at different developmental stages. In this regard, it is interesting to note that one of the first determinants of left-right asymmetry involves a flow in the node that has been shown to include lipoprotein vesicles containing Shh [81]. This occurs around E7.5, although signs of asymmetry are only apparent later in development [82, 83]. Therefore, the reduction in apolipoprotein-encoding genes could be involved in a restriction of the timing of this signaling. The Wnt-coreceptors Lrp5 and Lrp6 also belong to the family of lipoprotein receptors, thus adding the potential involvement of apolipoproteins in the differential regulation of Wnt signaling by modulating interactions between the Wnt molecules and their receptors.

### Transcription factor binding activity in the posterior epiblast

To assess how the modification of the chromatin accessibility profiles between E7.5 and E8.5 was reflected in the binding profiles of TFs known to be involved in developmental processes, we searched for TF footprints in our ATAC-seq datasets using HINT-ATAC [84]. We found several TFs with a significant difference in activity score between the two developmental stages (Fig 5A). At E7.5 we observed a higher activity score for TFs involved in pluripotency, like Pou5f1, Nanog and Sox2. The average ATAC-seq profiles around the binding sites of each of these TFs revealed that, although at a lower level and in a reduced number of regions, binding activity was still detected at E8.5 (Fig 5B-D). This might reflect a change in the functional profile of those factors as development proceeds. For instance, while Sox2 and Pou5f1 are required for pluripotency [23, 85], later in development they are involved in trunk elongation (Pou5f1) or in neural tube development (Sox2) [86–88]. At E8.5 the highest activity scores were provided by Cdx2, Cdx1, and several posterior Hox proteins (Fig 5A). Interestingly, their ATAC-seq profiles showed shallow footprints at E7.5 (Fig 5E-G), revealing that binding of these factors to their genomic targets mostly starts when the embryo engages in trunk development. These observations fit the genetic data showing that in the absence of Cdx activity, mouse embryos are truncated at the head to trunk transition [14,16,89], thus indicating that the functional requirement for these genes starts at this transition. Conversely, the binding profile of Tbxt (Brachyury), another of the main regulators of axial extension [13], was similar at E7.5 and E8.5 (Fig 5H). This suggests that Tbxt transcriptional activity might be similar at both developmental stages despite only being essential when the embryos enter trunk development [15, 89]. Alternatively, the Tbxt binding activity identified with the footprints at E7.5 might not be directly related to this protein but to Eomes, a T-box TF essential for endodermal and mesodermal development at early developmental stages [6] that shares DNA target sequence with Tbxt. Discerning between these possibilities will require evaluating Tbxt binding profiles in *Eomes* mutant embryos. Nfya and Nfyb were also found to have a higher activity score at E8.5 (Fig 5I). *Nfya* homozygous mutants exhibit early embryonic lethality [90], possibly related to the role of Nfya in zygotic genome activation [91]. Besides the role in early development, our data suggests that *Nfya* and *Nfyb* may also be important at the transition stage. Together, these results highlight a change in the main regulatory networks involved in each of these developmental stages, which is reflected by the activity levels of specific TFs.

**Figure 5.**
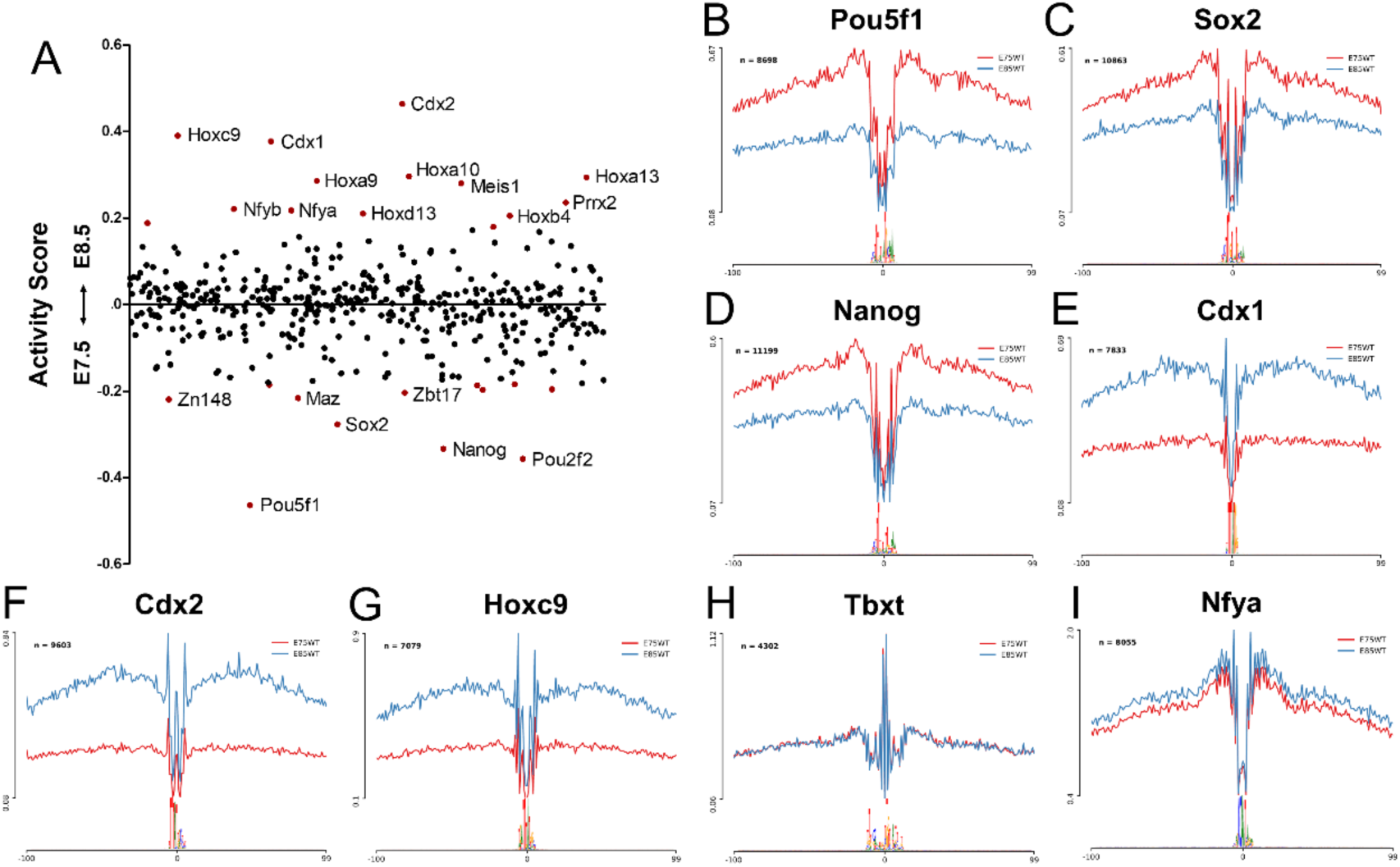
TF activity dynamics during the head to trunk transition. (A) Scatter plot of TF activity dynamics between E7.5 and E8.5. The y-axis represents the differences in TF binding activity. Each point represents a TF, points colored in red have significantly different activity scores (p-value < 0.05). Labelled points have a differential |Activity Score| > 0.2. (B-I) Average ATAC-seq profiles of Pou5f1 (B), Sox2 (C), Nanog (D), Cdx1 (E), Cdx2 (F), Hoxc9 (G), Tbxt (H) and Nfya (I) binding sites. Red profiles correspond to E7.5, blue profiles to E8.5, n indicates the number of binding sites used to calculate the average profiles.

### Testing a potential enhancer region of *Wnt5a*

From the 238 ATAC-seq peaks associated with differentially regulated genes, we focused on a region approximately 3.3kb upstream of *Wnt5a* transcriptional start site that becomes accessible at E8.5 (Fig 6A). This region is highly phylogenetically conserved among mammalian species [92], thus making it a candidate to regulate *Wnt5a* expression when the embryo engages in trunk development. We will refer to this region as CR1.

**Figure 6.**
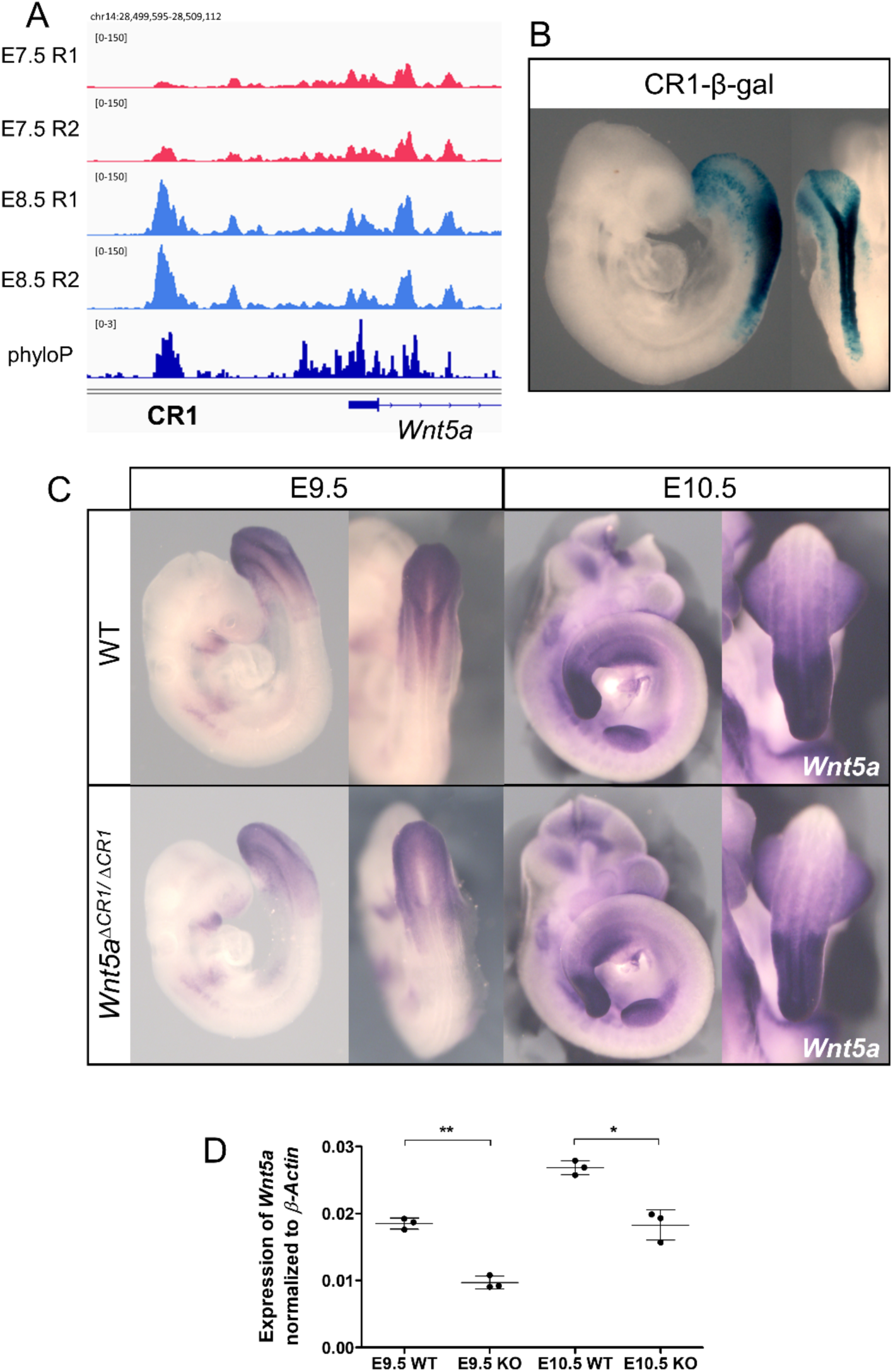
Characterization of Wnt5a enhancer, CR1. (A) ATAC-seq tracks showing accessibility profiles in the CR1 region. Phylogenetic conservation data (phyloP)[92] is shown in dark blue. (B) β-gal staining of *CR1-β-gal* transgenic embryo, also showing dorsal view of the caudal region. (C) Whole-mount *in situ* hybridization of wild type and *Wnt5a^ΔCR1/ΔCR1^* embryos at E9.5 and E10.5 using a probe for *Wnt5a*. RT-qPCR analysis of *Wnt5a* gene expression in wild type and *Wnt5a^ΔCR1/ΔCR1^* embryos at E9.5 and E10.5. *Wnt5a* expression is normalized to *β-Actin*. Error bars indicate the standard deviation; **, p-value < 0.01 and *, p-value < 0.05.

We first tested the regulatory potential of this putative enhancer using a reporter assay in mouse embryos. Transgenic embryos consistently displayed reporter expression in the posterior epiblast and emerging neural tube, a pattern closely resembling *Wnt5a* expression (Fig 6B). This pattern is consistent with CR1 involvement in *Wnt5a* activation in the progenitor region during the head to trunk transition.

To directly explore this hypothesis, we generated CR1 deletion mutants (*Wnt5a^ΔCR1^*). Whole-mount *in situ* hybridization suggested a reduction in *Wnt5a* expression levels in the caudal region of *Wnt5a^ΔCR1/ ΔCR1^* embryos at E9.5, and to a lesser extent at E10.5 (Fig 6C). This downregulation was confirmed by quantitative RT-PCR, at both stages (Fig 6D and S4 Table). These results suggest that, while the CR1 element participates in the regulation of *Wnt5a* expression in vivo, this regulation should also involve the activity of additional redundant enhancers that confer robustness to *Wnt5a* expression, able to keep a baseline *Wnt5a* transcription in *Wnt5a^ΔCR1/ ΔCR1^* mutants, thus allowing their full embryonic development. Despite the observed downregulation of *Wnt5a*, *Wnt5a^ΔCR1/ ΔCR1^* mutants developed normally, generating adult animals with no obvious phenotypic defects. This contrasts with *Wnt5a^-/-^* mutants, where loss of *Wnt5a* leads to perinatal lethality, with embryos showing an absence of tail and a shortening of the anterior-posterior axis [18].

### Impact of RA signaling on the transition from head to trunk development

Genetic analyses revealed a fundamental role of RA signaling for proper transition from head to trunk development [21]. We therefore tested the extent to which this is associated with changes in chromatin accessibility. We generated a *Raldh2* mutant strain by introducing in frame stop codons in the second exon (S2 Fig). Homozygous embryos for this strain showed a phenotype similar to that described for other previously described *Raldh2* mutants [21]. Comparison of the global accessibility profiles from the posterior epiblast region of E8.5 wild type and *Raldh2* mutant embryos revealed that only 120 peaks were differentially accessible between both conditions, including 54 regions with decreased and 66 regions with increased accessibility in the *Raldh2^-/-^* mutants (Fig 7A and S3 Table). Again, we observed that the differentially accessible peaks mapped mainly to intergenic regions (43%) (Fig 7B), indicating that they might represent regulatory elements.

**Figure 7.**
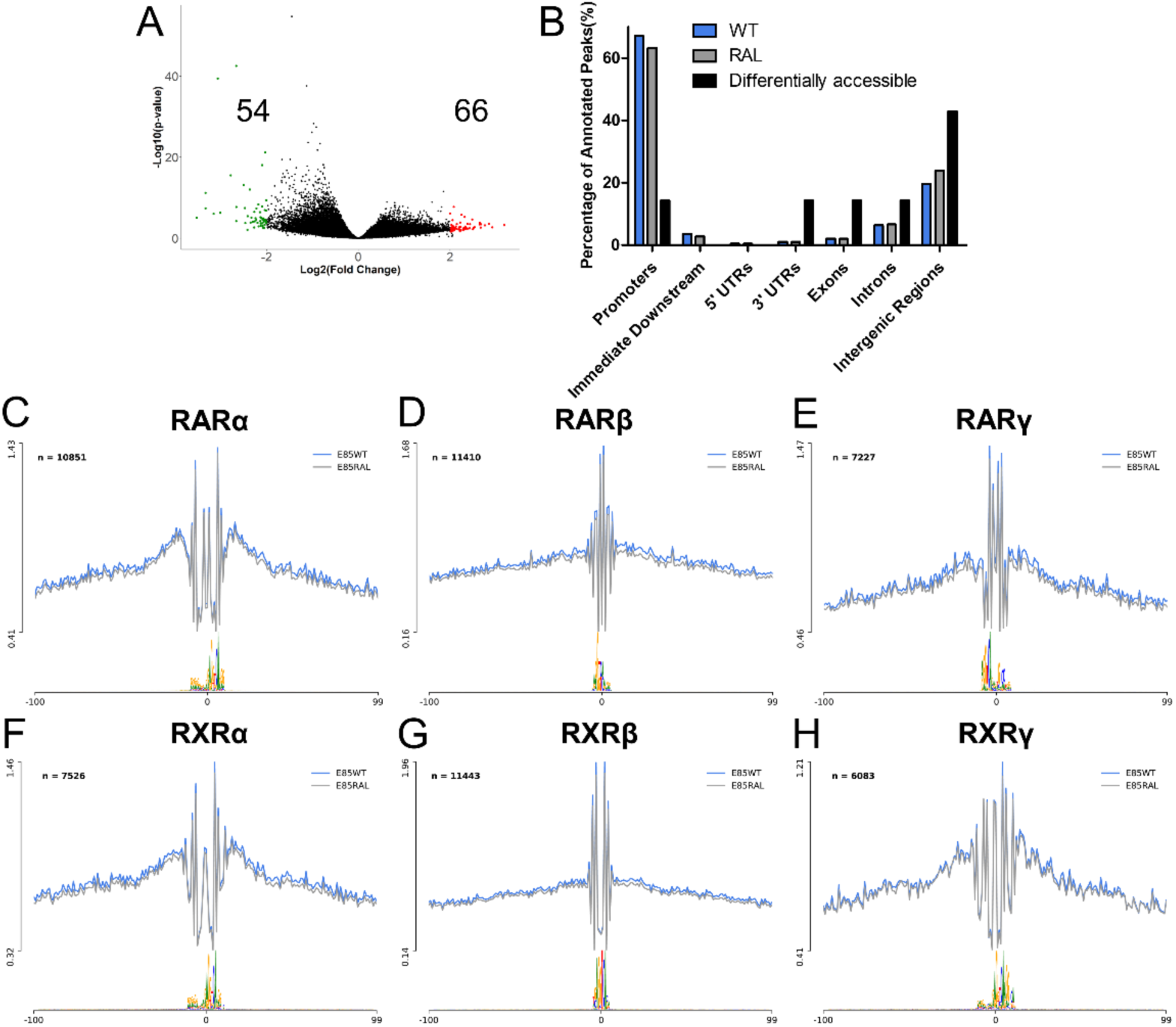
Impact of RA signaling in genome accessibility of the posterior epiblast. (A) Volcano plot of ATAC-seq peaks (*Raldh2^-/-^* vs WT) (|Log_2_(Fold Change)|>2 & p-value < 0.05). Significantly open regions in *Raldh2^-/-^* are red, closed regions in *Raldh2^-/-^* are green and non-significant in black. (B) Genomic distribution of ATAC-seq peaks identified in WT (blue), *Raldh2^-/-^* (gray) and distribution of only the differentially accessible peaks (black). (C-H) Average ATAC-seq profiles of RARα (C), RARβ (D), RARγ (E), RXRα (F), RXRβ (G) and RXRγ (H) binding sites. Blue profiles correspond to wild type, gray profiles to *Raldh2^-/-^*, n indicates the number of binding sites used to calculate the average profiles.

TF footprinting analyses showed no significant change in retinoic acid-and retinoid X receptor (RAR and RXR) binding activity in *Raldh2^-/-^* mutants (Fig 7C-H). This fits the notion that RA receptors are normally bound to retinoic acid response elements (RAREs) but kept inactive until bound by RA, eventually leading to recruitment of histone acetyltransferases and transcriptional activation [93]. Interestingly, only 12 of the regions that became differentially accessible contain RA receptor binding sites. Together, these observations suggest that RA activity at this developmental stage does not involve major changes in the genomic regions bound by RA receptors and that most of the differences in chromatin accessibility observed in the *Raldh2* mutants are not mediated by direct RA activity, most likely representing instead downstream effects of factors under direct RA regulation. The potential involvement of genes regulated by the 12 elements containing binding sequences for RA receptors in this or other RA-dependent regulatory processes will require direct experimental analyses.

### Evaluating potential *Nr2f2* enhancers for RA-mediated *Nr2f2* activation

From the regions that gained accessibility at E8.5 in a RA-dependent fashion we focused on a phylogenetically conserved region (we will refer to it as CR2) located within the same topologically associated domain (TAD) as *Nr2f2* (Fig 8A and 8B), a gene that has been shown to be under the control of RA signaling [94]. We selected CR2 for further analysis because when tested in transgenic reporter assays it reproduced to a large extent the *Nr2f2* expression pattern (Fig 8C), thus suggesting that it might be involved in the RA-dependent activation of *Nr2f2* expression. CR2 contains two distinct elements (CR2a and CR2b) (Fig 8A). Both elements also gave well-defined activity profiles when tested individually in transgenic reporter assays. *CR2a-β-gal* embryos displayed staining in the somites starting from the forelimb level, in rhombomere 5 and in the second branchial arch neural crest (Fig 8D – top panels). CR2b gave a much broader range of expression in the neural tube, including the whole hindbrain and the spinal cord, and in the neural crest migrating from the hindbrain into the branchial arches (Fig 8D – bottom panels). It also activated expression in the most anterior somites, where CR2a activity was not observed. Together, these staining patterns indicate that CR2 activity in the somites, branchial arches, and hindbrain might result from the combined CR2a and CR2b activities. However, the strong *CR2b-β-gal* reporter staining in the spinal cord contrasts with the absence of staining in the same region of CR2 reporter transgenics, suggesting that CR2a could block CR2b activity in this region.

**Figure 8.**
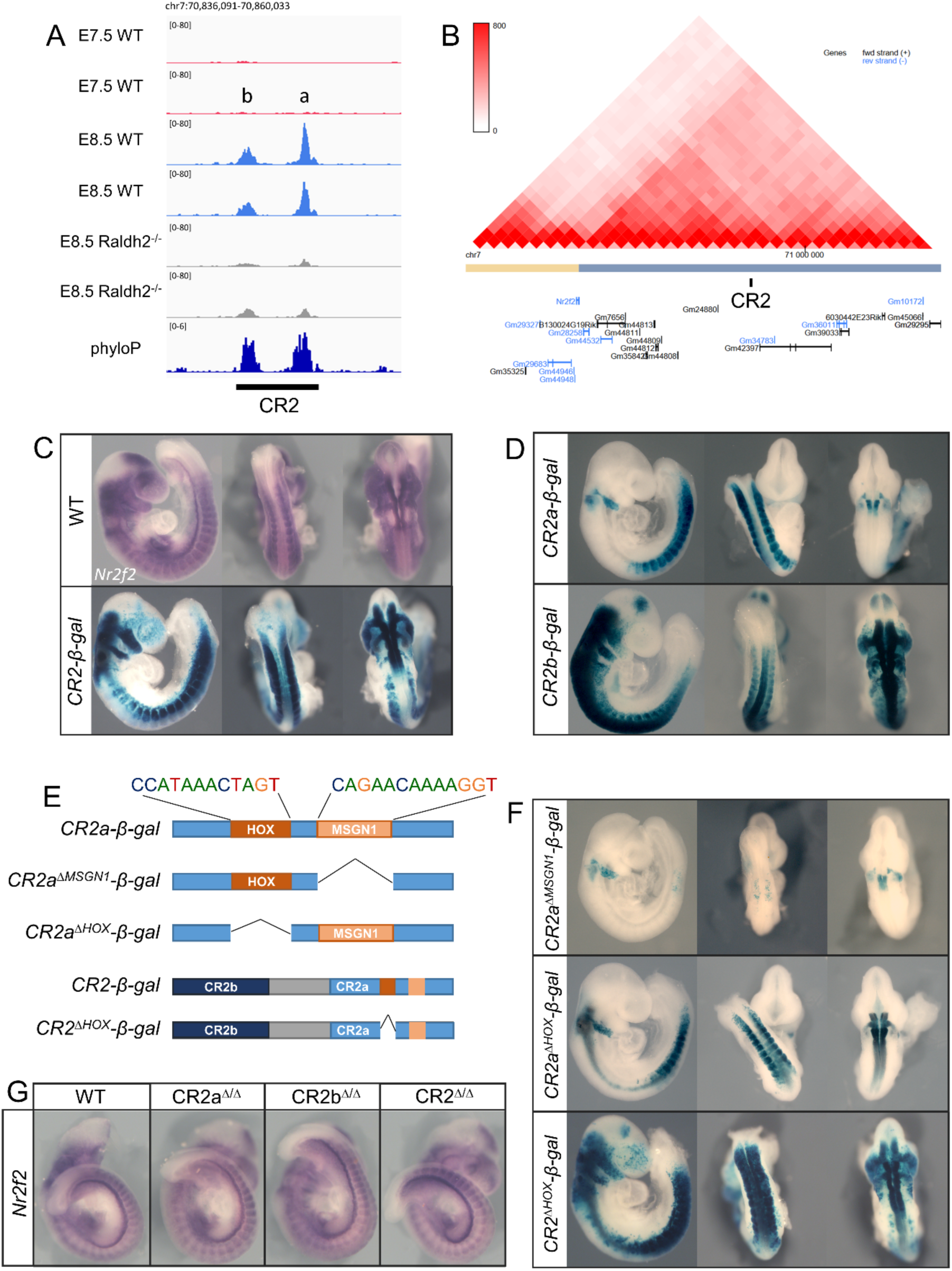
Characterization of the CR2 region. (A) ATAC-seq tracks showing accessibility profiles in the CR2 region. CR2 includes two peaks, *a* and *b*. Phylogenetic conservation data (phyloP)[92] is shown in dark blue. (B) Hi-C data from 3D genome browser [123] highlighting CR2 location in the same TAD as *Nr2f2*. (C) Comparison of *Nr2f2* expression pattern in wild type embryos by *in situ* hybridization (top panels) with β-gal reporter expression in *CR2-β-gal* transgenic embryos (bottom panels). (D) β-gal reporter expression in *CR2a-β-gal* (top panels) and *CR2b-β-gal* transgenic embryos (bottom panels). (C) schematic representation of generated transgenic reporters for CR2a and CR2 regions lacking the specified TF binding sites. (F) β-gal reporter expression in *CR2a^ΔMSGN1^*-β-gal (top panels), *CR2a^ΔHOX^*-β-gal (middle panels) and *CR2^ΔHOX^*-β-gal (bottom panels) transgenic embryos. (G) Whole-mount *in situ* hybridization of wild type, CR2a^Δ/Δ^, CR2b^Δ/Δ^ and CR2^Δ/Δ^ at E9.5 using a probe for Nr2f2.

To further analyze the mechanisms regulating CR2 enhancer activity and the interactions between CR2a and CR2b, we searched for the presence of TF binding sites within these elements with HINT-ATAC. We identified two TF footprints in CR2a, matching MSGN1 and HOX binding sites (Fig 8E). Given the important role of these TFs in embryonic development, we assessed their contribution to CR2a enhancer activity by generating transgenic reporters for the CR2a element lacking each of these features. Transgenic embryos generated with CR2a lacking the MSGN1 binding site (*CR2a^ΔMSGN1^*) lost almost completely reporter gene expression in the somites (Fig 8F – top panels), consistent with the known role of *Msgn1* as a regulator of paraxial mesoderm [95, 96]. Conversely, transgenic embryos of CR2a reporters lacking the HOX binding site (*CR2a^ΔHOX^*) did not affect somite expression, displaying instead extended reporter activity in the neural tube, including rhombomeres 3, 4 and 6 and the anterior spinal cord (Fig 8F – middle panels). This suggests a repressor rather than an activator role for Hox proteins in this enhancer, most particularly in the neural tube. We therefore tested whether the HOX binding site could also be involved in keeping CR2 inactive in the spinal cord by silencing CR2b activity in this embryonic region. Deletion of the HOX binding site from the CR2 reporter construct (*CR2^ΔHOX^*) resulted in a substantial activation of reporter activity in the neural tube, although not as extensive as the pattern obtained with CR2b (Fig 8D – bottom panels), indicating that it could indeed be part of the interaction mechanism between CR2a and CR2b.

We also identified binding sites for SMAD1 and SP5 in the CR2b element. Deletion of both sites resulted in the loss of reporter expression in most of the embryo, with some residual expression being detected in the hindbrain, neural crest and anterior spinal cord up until the trunk level (S3 Fig). Hence, the CR2a and CR2b elements are regulated by distinct sets of TFs, further allowing these regions to drive robust gene expression patterns despite possible fluctuations in upstream TF levels [97].

Together, the reporter assays indicate the existence of regulatory interactions between the CR2a and CR2b elements to achieve a pattern of activity resembling *Nr2f2* expression. CR2 thus represents a case in which enhancer interactions, both positive and negative, play an important role in fine tuning gene expression, contributing to the production of sharp boundaries in the expression domains, similarly to what has been previously reported for other systems [98–100]. Our results also suggest that the RA-dependent opening of CR2a and CR2b might expose these elements to become activated by factors involved in the development of trunk and hindbrain structures.

To directly assess CR2 function and its potential relevance for *Nr2f2* expression, we generated mouse strains containing deletions of CR2a, CR2b and CR2. Homozygous mutant animals for each of these strains developed to term and the adults had no obvious phenotypic alterations, already indicating that these mutants kept *Nr2f2* expression, at least to a level allowing normal development. We confirmed this by whole-mount *in situ* hybridization showing that the *Nr2f2* expression pattern in homozygous mutant embryos for any of the deleted CR2 regions were similar to that observed in wild type embryos (Fig 8G). This indicates that if CR2a and CR2b are indeed involved in *Nr2f2* expression as suggested by the reporter assays, other redundant enhancers might be present that ensure *Nr2f2* expression and prevent developmental arrest caused by the inactivation of this gene. A possible candidate for an enhancer able to maintain *Nr2f2* transcription in the absence of CR2 is the previously identified RARE in intron 1 of *Nr2f2* [94]. However, further studies will be required to validate the role of this RARE in the regulation of *Nr2f2* and whether it interacts functionally with CR2.

## Conclusion

Overall, this study provides a comprehensive analysis to gain insights into the mechanisms regulating the remarkable changes in tissue activity associated with the transition from head to trunk development, combining differential screening with bioinformatic treatment of the resulting data. The specific treatment of the DEG profiles with the identification of modules in the PPI network revealed changes in Wnt signaling, ubiquitination and the basic machinery of G-protein-mediated signal transduction that could engage in interactions resulting in a global functional output. In addition, our datasets can be used as a resource for future research not only to explore the role of other enhancer regions but also to delve into the mechanisms of gene regulatory networks involved in the head to trunk transition by combining it with gene knockout studies.

The high disparity in the number of changes in chromosomal accessibility and differentially expressed genes at the two developmental stages indicates that the control of the changes in gene expression might also be very complex. Consistent with this, the lack of obvious phenotypes upon deletion of regulatory elements that largely reproduce the expression profiles of the genes they might regulate, argues for the existence of a considerable degree of redundancies among regulatory mechanisms. This redundancy, which has been previously observed for other regulatory regions [101, 102], might confer robustness to developmental processes by providing protection against genetic and environmental perturbations [101,103–105]. Future studies testing the effects of CR1 or any of the CR2 deletions in a different genetic background or combined with a heterozygous inactivation of the *Wnt5a* or *Nr2f2* genes, respectively, or with other potential regulatory elements may reveal phenotypic traits normally suppressed by functional redundancy among enhancers.

## Methods

### Mice and Embryos

Mouse embryos were recovered by cesarean section at different developmental stages and processed accordingly for the distinct analysis described below.

*Raldh2* mutants, CR1, CR2, CR2a and CR2b deletion mice were generated by CRISPR/Cas9 [106] on the FVB/J background. *Raldh2* mutant mice were generated by introducing in frame stop codons into the second exon of the gene (S2 Fig). A sgRNA targeting the sequence AATGGCAGAACTCAGAGAGT was generated by in vitro transcription. Briefly, oligonucleotides Raldh2-gRNA-up and Raldh2-gRNA-down (Table 1) were annealed and cloned into the BssI sites of plasmid pgRNA-basic [107]. The sgRNA was transcribed from the resulting plasmid with the MEGAshortscript T7 Kit (Life Technologies) and purified with the MEGAclear Kit (Life Technologies). Cas9 mRNA was produced by in vitro transcription from the pT7-Cas9 plasmid [107] using the mMESSAGE mMACHINE T7 Ultra Kit (Life Technologies) and purified with the MEGAclear Kit (Life Technologies). The replacement ssDNA oligonucleotide containing three in frame stop codons followed by an EcoRI site (Raldh2-3X-Stop) (Table 1) was purchased from IDT. A mixture of 10 ng/μl of Cas9 mRNA, 10 ng/μl of the gRNA and 10 ng/μl of the Raldh2-3X-Stop oligonucleotide was injected into the pronuclei of fertilized oocytes of the FVB/J background, using standard procedures [108]. The mutant allele was detected by PCR using primers Raldh2-F and Raldh2-MUT-R (Table 2). Targeting was confirmed by direct sequencing. Deletion mutants for CR1, CR2, CR2a and CR2b were generated as previously described [109], using two gRNAs targeting the border of the sequence to be deleted and one ssDNA oligo bridging the two sides of the deletion (Table 1) to increase the edition efficiency. In these cases, each gRNA was generated by annealing the relevant Alt-R-CRISPR-Cas9 crRNA (targeting sequences in Table 1) with the Alt-R-CRISPR-Cas9 tracrRNA (all purchased from IDT). 1μM of each gRNA was incubated with 100 ng/μl of theCas9 protein and 10 ng/μl of the replacement DNA and microinjected into the pronucleus of fertilized mouse oocytes. Identification of deletion mutants was performed by PCR using the oligonucleotides specified in Table 2. Positive founders were crossed with wild type mice to generate F1 heterozygous mice that were then used to build the mutant lines. Homozygous mutants were then generated by heterozygous crosses. Mice and embryos were genotyped by PCR using primers specified on Table 2.

**Table 1.**
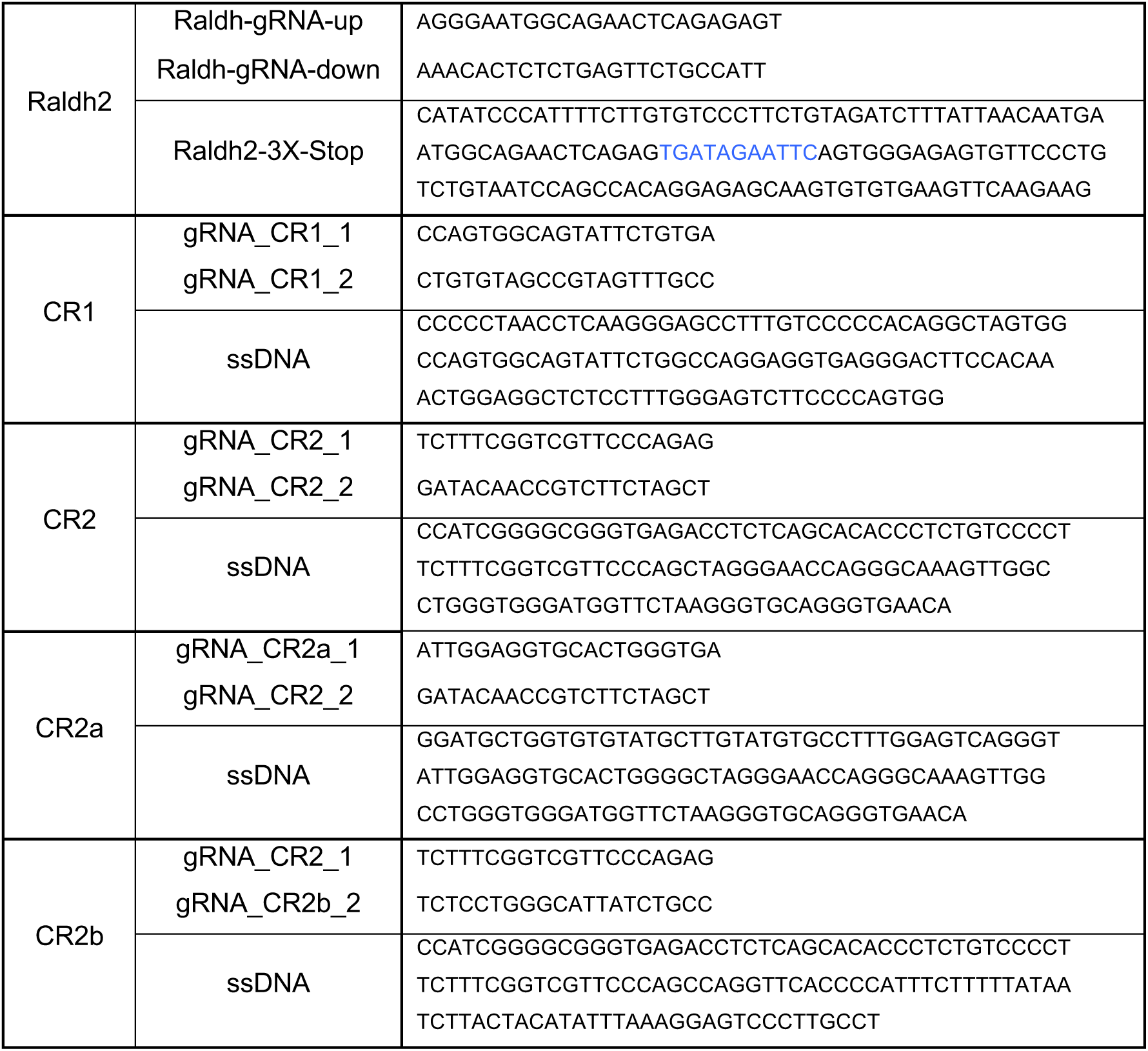
gRNA and ssDNA used for CRISPR/Cas9.

**Table 2.**
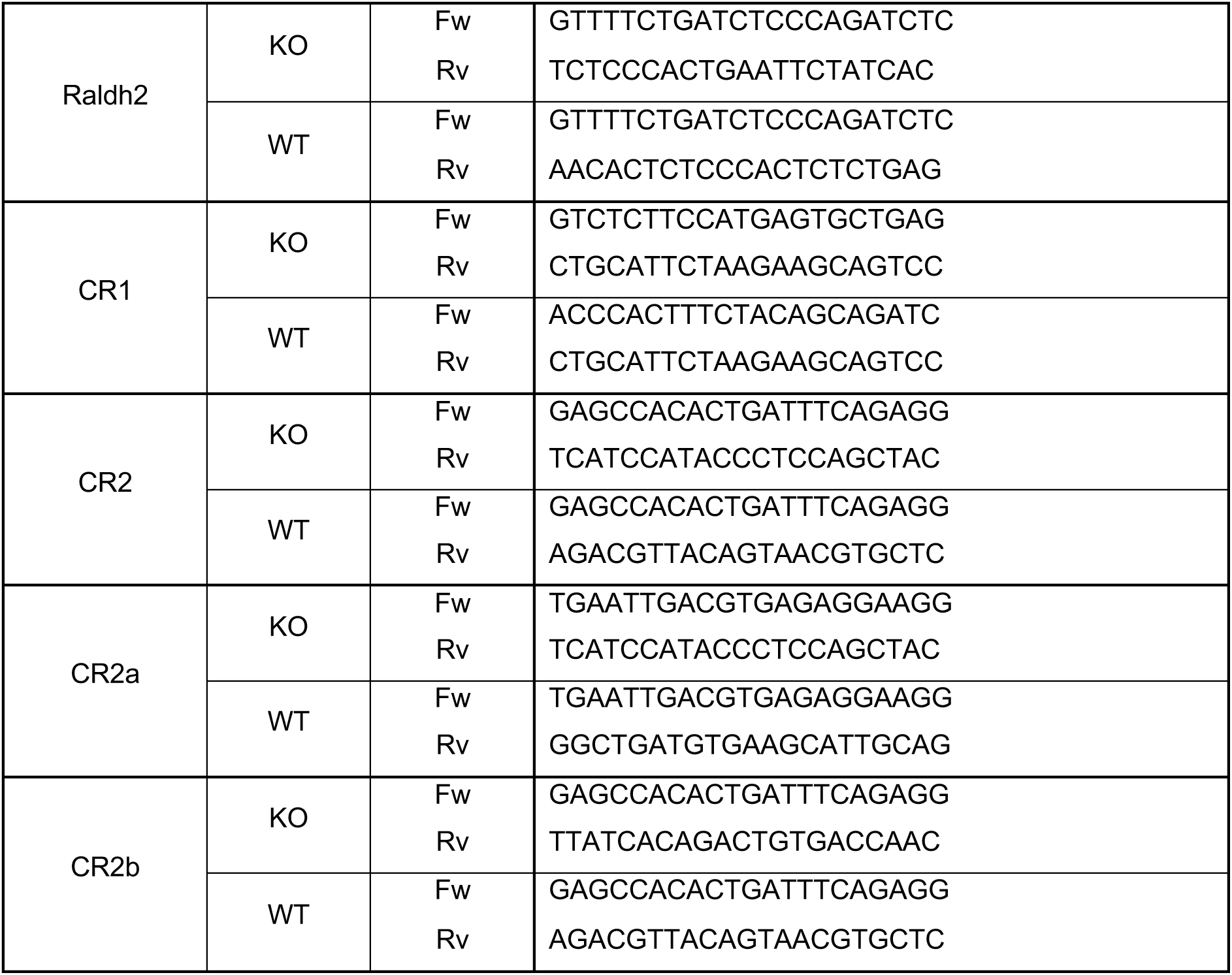
Primers used for genotyping.

All experiments conducted on animals followed the Portuguese (Portaria 1005/92) and European (Directive 2010/63/EU) legislations concerning housing, husbandry, and welfare. The project was reviewed and approved by the Ethics Committee of Instituto Gulbenkian de Ciência and by the Portuguese National Entity, Direcção Geral de Alimentação e Veterinária (license reference 014308).

### RNA-sequencing analysis

Posterior epiblasts of wild type mouse embryos at E7.5 and E8.5 were dissected and snap frozen. Total RNA was isolated from pooled samples with TRI Reagent following the manufacturer’s protocol. RNA samples were then resuspended in RNase-free water. RNA concentration and purity were determined on an AATI Fragment Analyzer (Agilent). RNA-seq from E7.5 and E8.5 tissues was performed using two separate biological replicates. Libraries were prepared from total RNA using the SMART-Seq2 protocol [110]. Sequencing was performed on Illumina NextSeq500, generating >25M single-end 75 base reads per sample. Reads were aligned to the reference mouse genome (mm10) using STAR [111]. Read count normalization and differential expression between samples was analyzed using DESeq2 [112]. RNA-seq data is available in the GEO accession database under the accession number GSE220246. Gene ontology enrichment analysis was performed using PANTHER [113, 114]. To assemble the PPI network, the DEG were filtered according to the following criteria: log of count per million (logCPM) >1; absolute log fold-change (logFC) >1; and false discovery rate (FDR) <0.05. All possible interactions between DEGs were retrieved from the STRING v11 protein-protein interactions database [35]. The mouse transcriptome network was then constructed from the set of expressed genes and their corresponding STRING PPI. We casted this network as a weighted graph, where edge weights (given by the STRING PPI scores) denote the probability of the connected genes interacting and thus jointly contributing to a specific function. To remove redundant edges and focus our attention on the most important interactions we extracted the (metric) backbone of the mouse transcriptome network [36]. The metric backbone is a subgraph that is sufficient to compute all shortest paths in the network, thus removing edges that break the triangle inequally (and are therefore redundant regarding the shortest paths). This network retains all metric edges and preserves all the nodes in the original network [36, 115]. We have previously used the metric backbone of transcriptome networks to identify biologically relevant genes and their interactions [37]. Network modules, i.e., structurally coherent structures in the transcriptome network backbone were identified using LowEnDe [38], an in-house developed algorithm based on the spectral decomposition of the adjacency matrix coupled with information theory to identify overlapping modules in weighted graphs. Importantly, in this method genes may participate in more than one module at the same time, reflecting the possible participation of genes in multiple cell functions.

### Embryo culture with Porcn inhibitor

Wild type E8.5 embryos were dissected, in cold GMEM (Sigma #G5154), keeping the yolk sac intact. Embryos were cultured in 60% Rat serum, 40% GMEM and Pen/Strep (Gibco #15070063). For embryos cultured with Porcn inhibitor, 500 nM of IWP-01 (MedChem express #HY-100853) was added as in [66], whereas for control embryos an equal volume of DMSO was added. Embryos were cultured for 24 hours in a rotator bottle culture apparatus (B.T.C. Engineering, Milton, Cambridge, UK) at 37°C, 65% O_2_. Three embryos were cultured per tube in 1,5 ml of media. Embryos were collected after 24hrs, dissected and fixed in 4% PFA at 4°C overnight. They were then processed for *in situ* hybridization, 2 embryos were stained per probe and condition, showing similar patterns. In addition, the structure of the neural tube was also assessed in the sections of embryos stained for other markers, showing highly reproducible patterns.

### ATAC-seq

Posterior epiblasts of mouse embryos were collected to 500 μl of cold M2 (Sigma #M7167), spun down to remove supernatant and incubated with 500 μl of Accutase (Sigma #A6964) for 30 min at 37°, with shaking at 600 rpm, to dissociate the tissue into single cells. ATAC-seq was performed as previously described [70], using two separate biological replicates for each condition. The amplified libraries were double-step size selected (0.5x followed by 1x) using SPRIselect (Beckman Coulter #B23317) according to manufacturer’s instructions. Pooled ATAC-seq libraries were sequenced on a NextSeq500 (Illumina) at 50M paired-end 75 base reads per sample.

### ATAC-seq Data Analysis

Fastq files were processed with GUAVA v1, following the recommended guidelines [116]. GUAVA enables pre-processing of raw sequencing reads, mapping of reads to a reference genome, peak calling and annotation, as well as differential analysis between samples. All genome browser tracks were captured using Integrative Genomics Viewer [117]. ATAC-seq data is available in the GEO accession database under accession number GSE220245.

ATAC-seq data was analyzed for TF footprints using HINT [84]. Replicates were merged to increase read depth and processed with “rgt-hint footprinting” command. Footprint motifs were matched to HOCOMOCO database [118] with “rgt-motifanalysis matching” and then further assessed for differential motif occupancy with the “rgt-hint differential” command.

### β-Galactosidase Transgenics

For reporter analyses, candidate regions identified by ATAC-seq data were amplified by PCR from mouse genomic DNA (primers provided below, Table 3) and cloned upstream of a cassette containing the adenovirus 2 minimal late promoter, the β-galactosidase cDNA, and the polyadenylation signal from SV40 [119]. Transgenic mice were produced by pronuclear injection [108]. The β-galactosidase staining was performed as previously described [119].

**Table 3.**
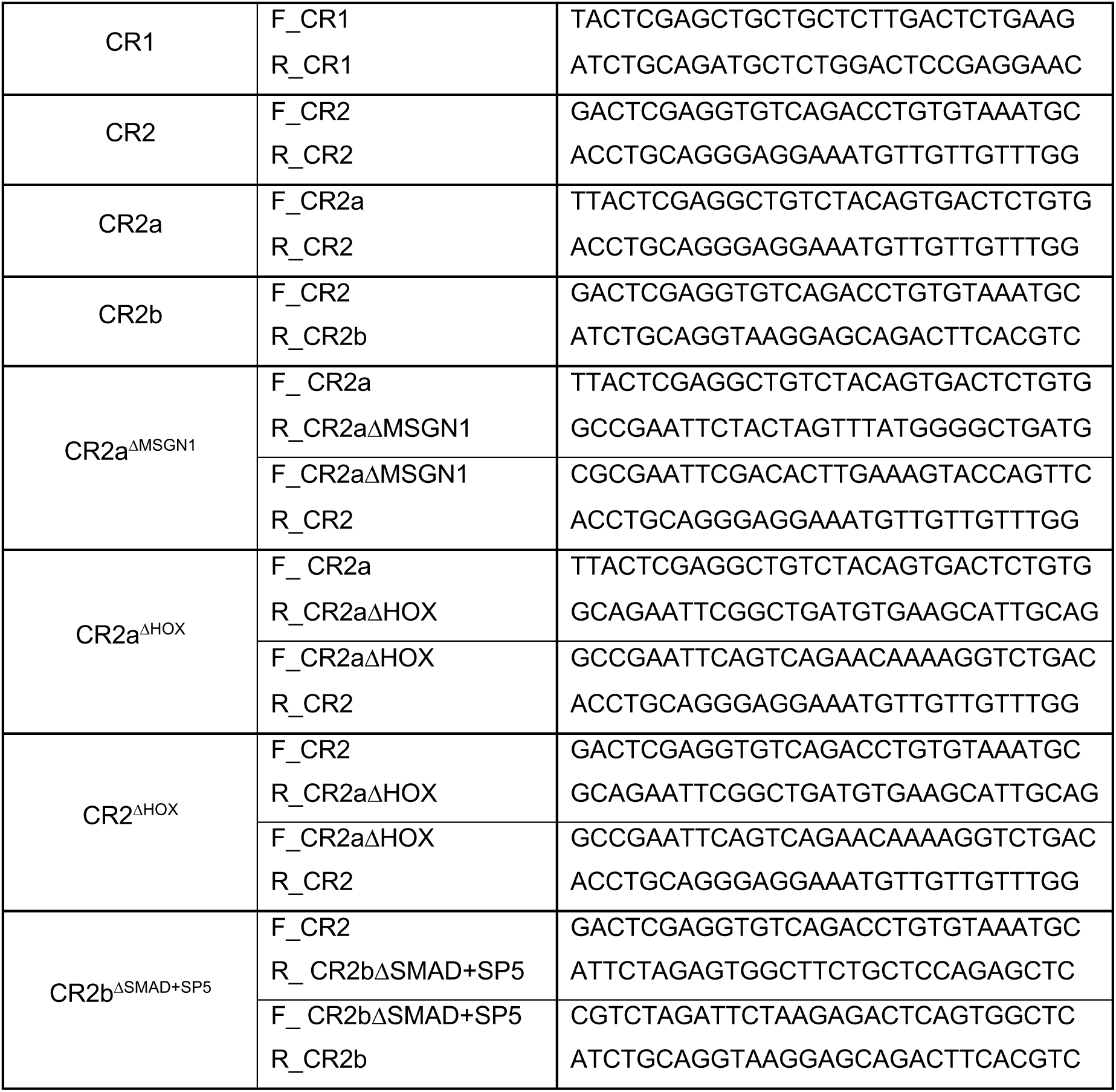
Primers used to amplify candidate regions for β-Galactosidase assays.

### Whole-mount *in situ* hybridization

Whole-mount in situ hybridization was performed as previously described [120] using digoxigenin-labeled RNA antisense probes. For the genetically modified embryos and their wild type controls, at least 3 embryos were stained per probe and genotype, showing highly reproducible patterns. RNA probes have been previously described: *Msgn1* [120]; *Sox2*, *Tbxt* and *Uncx4.1* [87]; *Shh* and *Cdx2* [119]; *Wnt5a* [121] and Fgf4 [122]. A probe for *Nr2f2* was prepared by amplifying a cDNA fragment by RT-PCR (primers provided below, Table 4) from total RNA isolated from E9.5 embryos with Tri Reagent (Sigma #93289) according to the manufacturer’s protocol, and cloning it into pKS-bluescript.

**Table 4.**
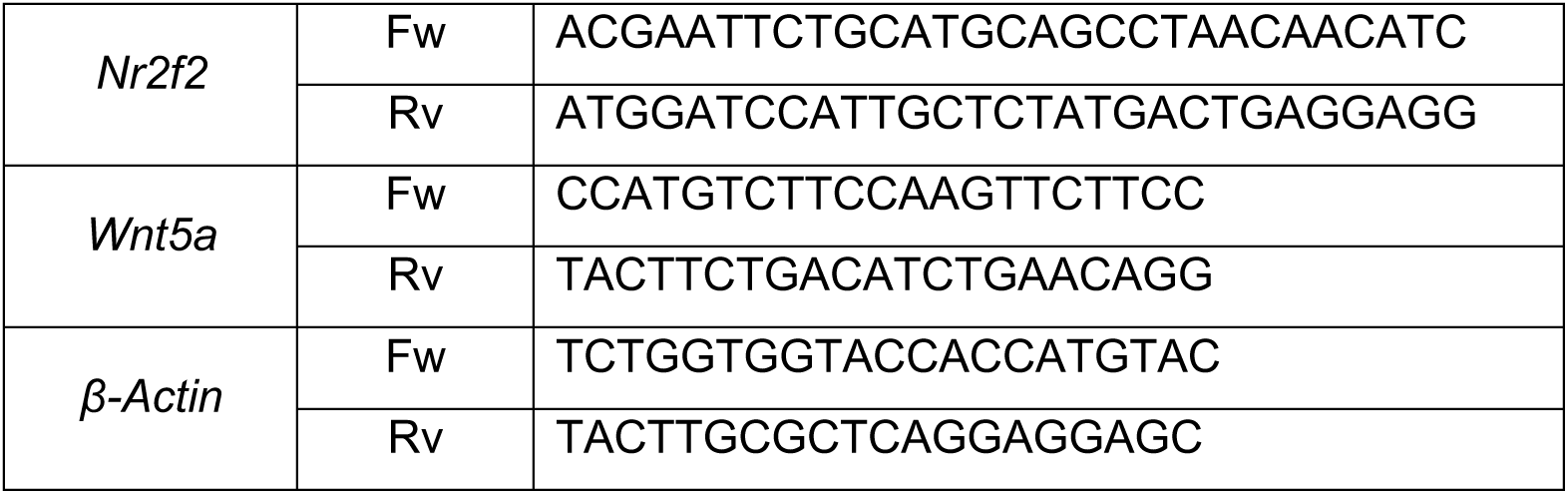
Primers used to amplify *in situ* probes and used in RT-qPCR.

### RT-qPCR

Total RNA was extracted from the caudal region of wild type and *Wnt5a^ΔCR1/ΔCR1^* embryos at E9.5 and E10.5 using Tri Reagent. 1 μg of RNA was used for reverse transcription into complementary DNA (cDNA) using NZY Reverse Transcriptase enzyme (NZYTech #MB124) and random hexamer mix (NZYTech #MB12901) following the manufacturer’s protocol. Real-time qPCR was performed in a QuantStudio 7 Flex real-time PCR system (Thermo Fisher) using iQ SYBR Green Supermix (Bio-rad #1708880) according to manufacturer’s instructions. Primers used are listed in Table 4. Quantification was determined using the standard curve method, and expression levels normalized to *β-Actin*. Statistical significance was assessed using paired t-test, **p<0.01 and *p<0.05.

## Supporting Information

S1 Table. Complete differential analysis results of RNA-seq datasets.

S2 Table. Complete differential analysis results of ATAC-seq E7.5 and E8.5 wild type datasets.

S3 Table. Complete differential analysis results of ATAC-seq wild type and *Raldh2^-/-^* datasets.

S4 Table. RT-qPCR data values of *Wnt5a* expression normalized to *β-Actin*.

S1 Network. Protein-protein interaction network generated with the differentially expressed genes.

## Acknowledgments

We thank the IGC animal house and Genomics Facilities for their expert services, advice and assistance. We also acknowledge André Dias for the help with the *Raldh2* colony, Anastasiia Lozovska for the help with embryo culture and RT-qPCR data analysis, and all members of the Mallo laboratory for helpful discussions and comments throughout the course of the project.

## Supplementary Figures

**S1 Fig.**
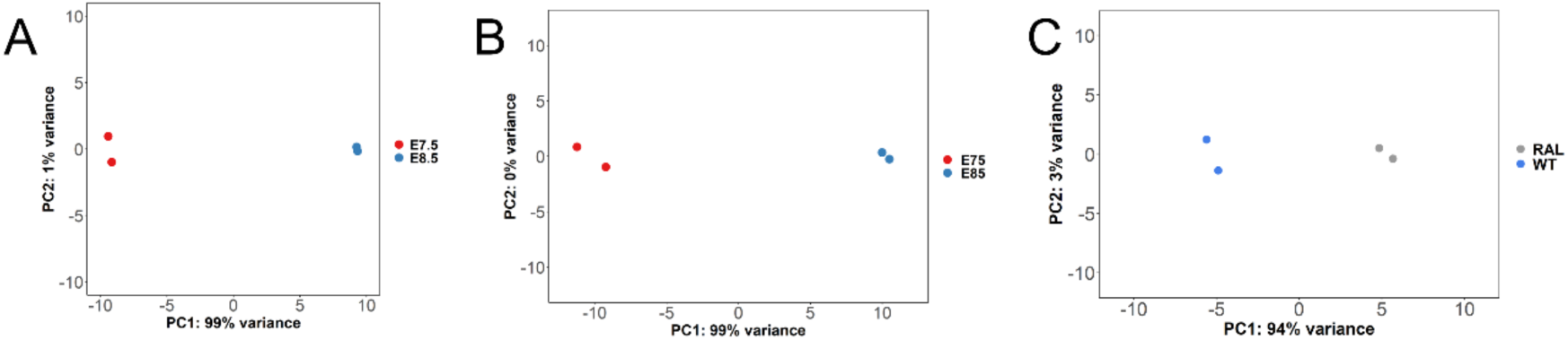
PCA plots of all the genome wide analysis. (A) Principal Component Analysis of RNA-seq data from E7.5 (red) and E8.5 (blue). (B) Principal Component Analysis of ATAC-seq data from E7.5 (red) and E8.5 (blue). (C) Principal Component Analysis of ATAC-seq data from WT (blue) and *Raldh2^-/-^* (grey).

**S2 Fig.**
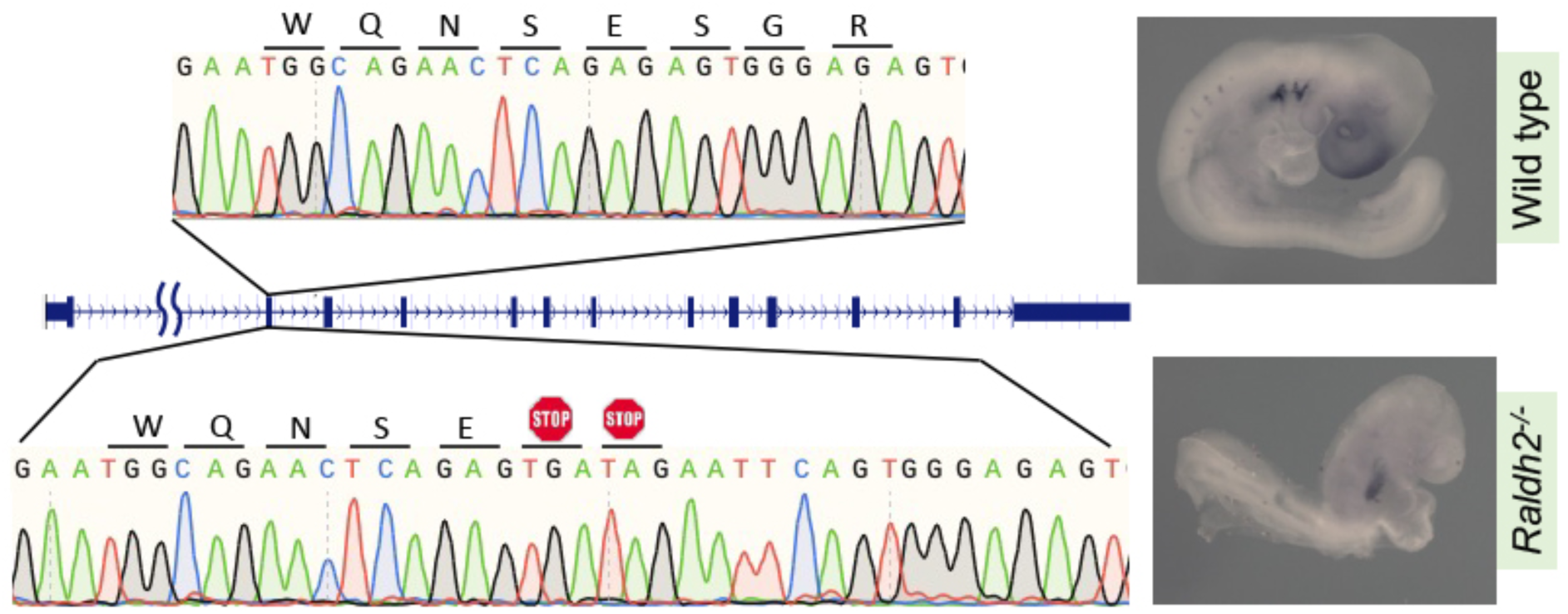
Characterization of *Raldh2^-/-^* mutants. Sequencing profiles of wild type and *Raldh2^-/-^* mutants, generated by introducing in frame stop codons in the second exon. Whole-mount in situ hybridization of wild type and *Raldh2^-/-^* mutant embryos at E9.5 using a probe for *Fgf4*.

**S3 Fig.**
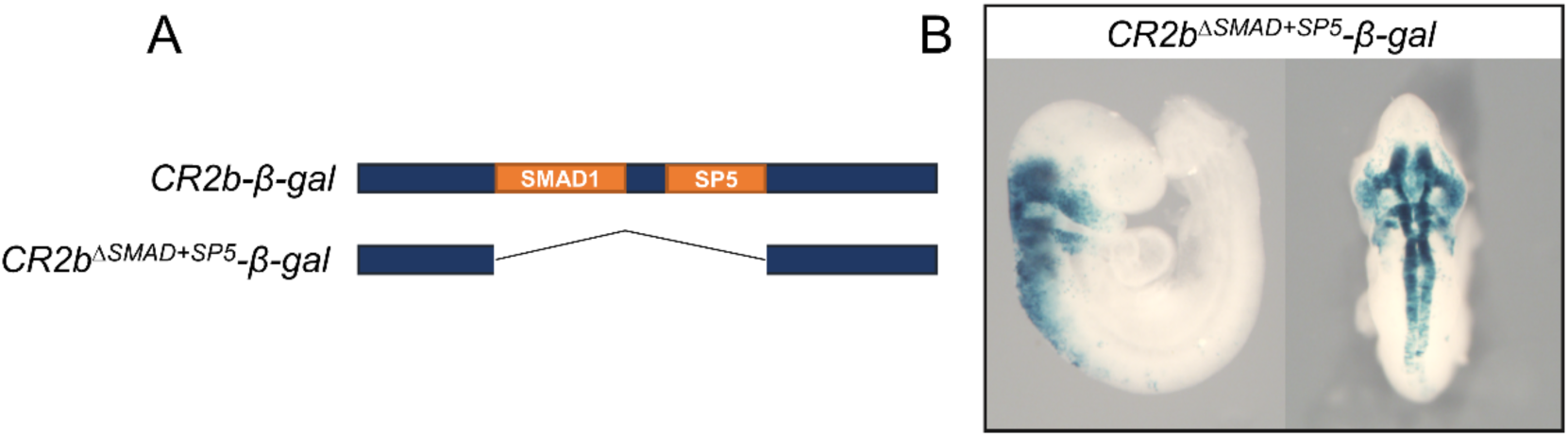
TF activity in the regulation of the CR2b element. (A) Schematic representation of generated transgenic reporters for CR2b lacking the specified TF binding sites. (B) *β-gal* reporter expression in *CR2b^ΔSMAD1+SP5^-β-gal* transgenics, lateral and dorsal views.

## Notes

### Competing Interest Statement

The authors have declared no competing interest.

## References

1. Stern CD, Charite J, Deschamps J, Duboule D, Durston AJ, Kmita M, et al. Head-tail patterning of the vertebrate embryo: one, two or many unresolved problems? Int J Dev Biol. 2006;50: 3– 15. doi:10.1387/ijdb.052095cs

2. Aires R, Dias A, Mallo M. Deconstructing the molecular mechanisms shaping the vertebrate body plan. Current Opinion in Cell Biology. 2018;55: 81–86. doi:10.1016/j.ceb.2018.05.009

3. Wilson V, Olivera-Martinez I, Storey KG. Stem cells, signals and vertebrate body axis extension. Development. 2009;136: 1591–1604. doi:10.1242/dev.021246

4. Tam PP, Behringer RR. Mouse gastrulation: the formation of a mammalian body plan. Mech Dev. 1997;68: 3–25. doi:10.1016/s0925-4773(97)00123-8

5. Bardot ES, Hadjantonakis A-K. Mouse gastrulation: Coordination of tissue patterning, specification and diversification of cell fate. Mechanisms of Development. 2020;163: 103617. doi:10.1016/j.mod.2020.103617

6. Arnold SJ, Hofmann UK, Bikoff EK, Robertson EJ. Pivotal roles for eomesodermin during axis formation, epithelium-to-mesenchyme transition and endoderm specification in the mouse. Development. 2008;135: 501–511. doi:10.1242/dev.014357

7. Zhao R, Watt AJ, Battle MA, Li J, Bondow BJ, Duncan SA. Loss of both GATA4 and GATA6 blocks cardiac myocyte differentiation and results in acardia in mice. Dev Biol. 2008;317: 614– 619. doi:10.1016/j.ydbio.2008.03.013

8. Steventon B, Martinez-Arias A. Evo-engineering and the cellular and molecular origins of the vertebrate spinal cord. Dev Biol. 2017;432: 3–13. doi:10.1016/j.ydbio.2017.01.021

9. Wymeersch FJ, Wilson V, Tsakiridis A. Understanding axial progenitor biology in vivo and in vitro. Development. 2021;148: dev180612. doi:10.1242/dev.180612

10. Binagui-Casas A, Dias A, Guillot C, Metzis V, Saunders D. Building consensus in neuromesodermal research: Current advances and future biomedical perspectives. Curr Opin Cell Biol. 2021;73: 133–140. doi:10.1016/j.ceb.2021.08.003

11. Ferretti E, Hadjantonakis A-K. Mesoderm specification and diversification: from single cells to emergent tissues. Curr Opin Cell Biol. 2019;61: 110–116. doi:10.1016/j.ceb.2019.07.012

12. Wymeersch FJ, Huang Y, Blin G, Cambray N, Wilkie R, Wong FC, et al. Position-dependent plasticity of distinct progenitor types in the primitive streak. Bronner ME, editor. eLife. 2016;5: e10042. doi:10.7554/eLife.10042

13. Herrmann BG, Labeit S, Poustka A, King TR, Lehrach H. Cloning of the T gene required in mesoderm formation in the mouse. Nature. 1990;343: 617–622. doi:10.1038/343617a0

14. Savory JGA, Mansfield M, Rijli FM, Lohnes D. Cdx mediates neural tube closure through transcriptional regulation of the planar cell polarity gene Ptk7. Development. 2011;138: 1361–1370. doi:10.1242/dev.056622

15. Chesley P. Development of the short-tailed mutant in the house mouse. 1935. doi:10.1002/JEZ.1400700306

16. Chawengsaksophak K, James R, Hammond VE, Köntgen F, Beck F. Homeosis and intestinal tumours in Cdx2 mutant mice. Nature. 1997;386: 84–87. doi:10.1038/386084a0

17. Takada S, Stark KL, Shea MJ, Vassileva G, McMahon JA, McMahon AP. Wnt-3a regulates somite and tailbud formation in the mouse embryo. Genes Dev. 1994;8: 174–189. doi:10.1101/gad.8.2.174

18. Yamaguchi TP, Bradley A, McMahon AP, Jones S. A Wnt5a pathway underlies outgrowth of multiple structures in the vertebrate embryo. Development. 1999;126: 1211–1223. doi:10.1242/dev.126.6.1211

19. Andre P, Song H, Kim W, Kispert A, Yang Y. Wnt5a and Wnt11 regulate mammalian anterior-posterior axis elongation. Development. 2015;142: 1516–1527. doi:10.1242/dev.119065

20. Rhinn M, Dollé P. Retinoic acid signalling during development. Development. 2012;139: 843– 858. doi:10.1242/dev.065938

21. Niederreither K, Subbarayan V, Dollé P, Chambon P. Embryonic retinoic acid synthesis is essential for early mouse post-implantation development. Nat Genet. 1999;21: 444–448. doi:10.1038/7788

22. Zhao X, Sirbu IO, Mic FA, Molotkova N, Molotkov A, Kumar S, et al. Retinoic acid promotes limb induction through effects on body axis extension but is unnecessary for limb patterning. Curr Biol. 2009;19: 1050–1057. doi:10.1016/j.cub.2009.04.059

23. Nichols J, Zevnik B, Anastassiadis K, Niwa H, Klewe-Nebenius D, Chambers I, et al. Formation of pluripotent stem cells in the mammalian embryo depends on the POU transcription factor Oct4. Cell. 1998;95: 379–391. doi:10.1016/s0092-8674(00)81769-9

24. Mitsui K, Tokuzawa Y, Itoh H, Segawa K, Murakami M, Takahashi K, et al. The homeoprotein Nanog is required for maintenance of pluripotency in mouse epiblast and ES cells. Cell. 2003;113: 631–642. doi:10.1016/s0092-8674(03)00393-3

25. Conlon FL, Barth KS, Robertson EJ. A novel retrovirally induced embryonic lethal mutation in the mouse: assessment of the developmental fate of embryonic stem cells homozygous for the 413.d proviral integration. Development. 1991;111: 969–981. doi:10.1242/dev.111.4.969

26. Ding J, Yang L, Yan YT, Chen A, Desai N, Wynshaw-Boris A, et al. Cripto is required for correct orientation of the anterior-posterior axis in the mouse embryo. Nature. 1998;395: 702–707. doi:10.1038/27215

27. Mitiku N, Baker JC. Genomic Analysis of Gastrulation and Organogenesis in the Mouse. Developmental Cell. 2007;13: 897–907. doi:10.1016/j.devcel.2007.10.004

28. Jaffe LF. The role of ionic currents in establishing developmental pattern. Philos Trans R Soc Lond B Biol Sci. 1981;295: 553–566. doi:10.1098/rstb.1981.0160

29. Levin M. Bioelectric signaling: Reprogrammable circuits underlying embryogenesis, regeneration, and cancer. Cell. 2021;184: 1971–1989. doi:10.1016/j.cell.2021.02.034

30. Levin M, Thorlin T, Robinson KR, Nogi T, Mercola M. Asymmetries in H+/K+-ATPase and cell membrane potentials comprise a very early step in left-right patterning. Cell. 2002;111: 77– 89. doi:10.1016/s0092-8674(02)00939-x

31. Raya Á, Kawakami Y, Rodríguez-Esteban C, Ibañes M, Rasskin-Gutman D, Rodríguez-León J, et al. Notch activity acts as a sensor for extracellular calcium during vertebrate left–right determination. Nature. 2004;427: 121–128. doi:10.1038/nature02190

32. Schwab A, Fabian A, Hanley PJ, Stock C. Role of ion channels and transporters in cell migration. Physiol Rev. 2012;92: 1865–1913. doi:10.1152/physrev.00018.2011

33. Becchetti A, Munaron L, Arcangeli A. The role of ion channels and transporters in cell proliferation and cancer. Front Physiol. 2013;4: 312. doi:10.3389/fphys.2013.00312

34. Zhang R, Kang R, Klionsky DJ, Tang D. Ion Channels and Transporters in Autophagy. Autophagy. 2022;18: 4–23. doi:10.1080/15548627.2021.1885147

35. Szklarczyk D, Gable AL, Lyon D, Junge A, Wyder S, Huerta-Cepas J, et al. STRING v11: protein-protein association networks with increased coverage, supporting functional discovery in genome-wide experimental datasets. Nucleic Acids Res. 2019;47: D607–D613. doi:10.1093/nar/gky1131

36. Simas T, Correia RB, Rocha LM. The distance backbone of complex networks. Journal of Complex Networks. 2021;9: cnab021. doi:10.1093/comnet/cnab021

37. Correia RB, Almeida JM, Wyrwoll MJ, Julca I, Sobral D, Misra CS, et al. The conserved transcriptional program of metazoan male germ cells uncovers ancient origins of human infertility. bioRxiv; 2022. p. 2022.03.02.482557. doi:10.1101/2022.03.02.482557

38. Correia RB, Costa PN, Rocha LM. Extraction of overlapping modules in networks via spectral methods and information theory. Madrid, Spain; 2020.

39. Zheng N, Shabek N. Ubiquitin Ligases: Structure, Function, and Regulation. Annu Rev Biochem. 2017;86: 129–157. doi:10.1146/annurev-biochem-060815-014922

40. Koo B-K, Spit M, Jordens I, Low TY, Stange DE, van de Wetering M, et al. Tumour suppressor RNF43 is a stem-cell E3 ligase that induces endocytosis of Wnt receptors. Nature. 2012;488: 665–669. doi:10.1038/nature11308

41. Abrami L, Kunz B, Iacovache I, van der Goot FG. Palmitoylation and ubiquitination regulate exit of the Wnt signaling protein LRP6 from the endoplasmic reticulum. Proceedings of the National Academy of Sciences. 2008;105: 5384–5389. doi:10.1073/pnas.0710389105

42. Angers S, Thorpe CJ, Biechele TL, Goldenberg SJ, Zheng N, MacCoss MJ, et al. The KLHL12– Cullin-3 ubiquitin ligase negatively regulates the Wnt–β-catenin pathway by targeting Dishevelled for degradation. Nat Cell Biol. 2006;8: 348–357. doi:10.1038/ncb1381

43. Choi J, Park SY, Costantini F, Jho E, Joo C-K. Adenomatous Polyposis Coli Is Down-regulated by the Ubiquitin-Proteasome Pathway in a Process Facilitated by Axin*. Journal of Biological Chemistry. 2004;279: 49188–49198. doi:10.1074/jbc.M404655200

44. Kim S, Jho E. The Protein Stability of Axin, a Negative Regulator of Wnt Signaling, Is Regulated by Smad Ubiquitination Regulatory Factor 2 (Smurf2)*. Journal of Biological Chemistry. 2010;285: 36420–36426. doi:10.1074/jbc.M110.137471

45. Hart M, Concordet J-P, Lassot I, Albert I, del los Santos R, Durand H, et al. The F-box protein β-TrCP associates with phosphorylated β-catenin and regulates its activity in the cell. Current Biology. 1999;9: 207–211. doi:10.1016/S0960-9822(99)80091-8

46. Kitagawa M, Hatakeyama S, Shirane M, Matsumoto M, Ishida N, Hattori K, et al. An F-box protein, FWD1, mediates ubiquitin-dependent proteolysis of beta-catenin. EMBO J. 1999;18: 2401–2410. doi:10.1093/emboj/18.9.2401

47. Liu P, Wakamiya M, Shea MJ, Albrecht U, Behringer RR, Bradley A. Requirement for Wnt3 in vertebrate axis formation. Nat Genet. 1999;22: 361–365. doi:10.1038/11932

48. Winston JT, Strack P, Beer-Romero P, Chu CY, Elledge SJ, Harper JW. The SCFβ-TRCP–ubiquitin ligase complex associates specifically with phosphorylated destruction motifs in IκBα and β-catenin and stimulates IκBα ubiquitination in vitro. Genes Dev. 1999;13: 270–283.

49. Dorrello NV, Peschiaroli A, Guardavaccaro D, Colburn NH, Sherman NE, Pagano M. S6K1-and ßTRCP-Mediated Degradation of PDCD4 Promotes Protein Translation and Cell Growth. Science. 2006;314: 467–471. doi:10.1126/science.1130276

50. Müerköster S, Arlt A, Sipos B, Witt M, Großmann M, Klöppel G, et al. Increased Expression of the E3-Ubiquitin Ligase Receptor Subunit βTRCP1 Relates to Constitutive Nuclear Factor-κB Activation and Chemoresistance in Pancreatic Carcinoma Cells. Cancer Research. 2005;65: 1316–1324. doi:10.1158/0008-5472.CAN-04-1626

51. Schiffrin EL. The Endothelin System. In: Lip GYH, Hall JE, editors. Comprehensive Hypertension. Philadelphia: Mosby; 2007. pp. 317–323. doi:10.1016/B978-0-323-03961-1.50031-3

52. Liu Z, Yan S, Wang J, Xu Y, Wang Y, Zhang S, et al. Endothelial adenosine A2a receptor-mediated glycolysis is essential for pathological retinal angiogenesis. Nat Commun. 2017;8: 584. doi:10.1038/s41467-017-00551-2

53. Chigurupati S, Kulkarni T, Thomas S, Shah G. Calcitonin stimulates multiple stages of angiogenesis by directly acting on endothelial cells. Cancer Res. 2005;65: 8519–8529. doi:10.1158/0008-5472.CAN-05-0848

54. Wong C, Mahapatra NR, Chitbangonsyn S, Mahboubi P, Mahata M, Mahata SK, et al. The angiotensin II receptor (Agtr1a): functional regulatory polymorphisms in a locus genetically linked to blood pressure variation in the mouse. Physiological Genomics. 2003;14: 83–93. doi:10.1152/physiolgenomics.00162.2002

55. Ara T, Nakamura Y, Egawa T, Sugiyama T, Abe K, Kishimoto T, et al. Impaired colonization of the gonads by primordial germ cells in mice lacking a chemokine, stromal cell-derived factor-1 (SDF-1). Proc Natl Acad Sci U S A. 2003;100: 5319–5323. doi:10.1073/pnas.0730719100

56. Abedini A, Sayed C, Carter LE, Boerboom D, Vanderhyden BC. Non-canonical WNT5a regulates Epithelial-to-Mesenchymal Transition in the mouse ovarian surface epithelium. Sci Rep. 2020;10: 9695. doi:10.1038/s41598-020-66559-9

57. Pandur P, Läsche M, Eisenberg LM, Kühl M. Wnt-11 activation of a non-canonical Wnt signalling pathway is required for cardiogenesis. Nature. 2002;418: 636–641. doi:10.1038/nature00921

58. Slusarski DC, Corces VG, Moon RT. Interaction of Wnt and a Frizzled homologue triggers G-protein-linked phosphatidylinositol signalling. Nature. 1997;390: 410–413. doi:10.1038/37138

59. Ho H-YH, Susman MW, Bikoff JB, Ryu YK, Jonas AM, Hu L, et al. Wnt5a–Ror–Dishevelled signaling constitutes a core developmental pathway that controls tissue morphogenesis. Proceedings of the National Academy of Sciences. 2012;109: 4044–4051. doi:10.1073/pnas.1200421109

60. Brinkmann E-M, Mattes B, Kumar R, Hagemann AIH, Gradl D, Scholpp S, et al. Secreted Frizzled-related Protein 2 (sFRP2) Redirects Non-canonical Wnt Signaling from Fz7 to Ror2 during Vertebrate Gastrulation*. Journal of Biological Chemistry. 2016;291: 13730–13742. doi:10.1074/jbc.M116.733766

61. Satoh W, Gotoh T, Tsunematsu Y, Aizawa S, Shimono A. Sfrp1 and Sfrp2 regulate anteroposterior axis elongation and somite segmentation during mouse embryogenesis. Development. 2006;133: 989–999. doi:10.1242/dev.02274

62. Qian L, Mahaffey JP, Alcorn HL, Anderson KV. Tissue-specific roles of Axin2 in the inhibition and activation of Wnt signaling in the mouse embryo. Proceedings of the National Academy of Sciences. 2011;108: 8692–8697. doi:10.1073/pnas.1100328108

63. Takada R, Satomi Y, Kurata T, Ueno N, Norioka S, Kondoh H, et al. Monounsaturated Fatty Acid Modification of Wnt Protein: Its Role in Wnt Secretion. Developmental Cell. 2006;11: 791–801. doi:10.1016/j.devcel.2006.10.003

64. Rios-Esteves J, Haugen B, Resh MD. Identification of key residues and regions important for porcupine-mediated Wnt acylation. J Biol Chem. 2014;289: 17009–17019. doi:10.1074/jbc.M114.561209

65. Biechele S, Cox BJ, Rossant J. Porcupine homolog is required for canonical Wnt signaling and gastrulation in mouse embryos. Dev Biol. 2011;355: 275–285. doi:10.1016/j.ydbio.2011.04.029

66. Galli LM, Burrus LW. Differential Palmit(e)oylation of Wnt1 on C93 and S224 Residues Has Overlapping and Distinct Consequences. PLOS ONE. 2011;6: e26636. doi:10.1371/journal.pone.0026636

67. Ikeya M, Lee SMK, Johnson JE, McMahon AP, Takada S. Wnt signalling required for expansion of neural crest and CNS progenitors. Nature. 1997;389: 966–970. doi:10.1038/40146

68. Rao DM, Shackleford MT, Bordeaux EK, Sottnik JL, Ferguson RL, Yamamoto TM, et al. Wnt family member 4 (WNT4) and WNT3A activate cell-autonomous Wnt signaling independent of porcupine O-acyltransferase or Wnt secretion. J Biol Chem. 2019;294: 19950–19966. doi:10.1074/jbc.RA119.009615

69. Yoshikawa Y, Fujimori T, McMahon AP, Takada S. Evidence that absence of Wnt-3a signaling promotes neuralization instead of paraxial mesoderm development in the mouse. Dev Biol. 1997;183: 234–242. doi:10.1006/dbio.1997.8502

70. Corces MR, Trevino AE, Hamilton EG, Greenside PG, Sinnott-Armstrong NA, Vesuna S, et al. An improved ATAC-seq protocol reduces background and enables interrogation of frozen tissues. Nat Methods. 2017;14: 959–962. doi:10.1038/nmeth.4396

71. Beccari L, Yakushiji-Kaminatsui N, Woltering JM, Necsulea A, Lonfat N, Rodríguez-Carballo E, et al. A role for HOX13 proteins in the regulatory switch between TADs at the HoxD locus. Genes Dev. 2016;30: 1172–1186. doi:10.1101/gad.281055.116

72. Okazawa H, Okamoto K, Ishino F, Ishino-Kaneko T, Takeda S, Toyoda Y, et al. The oct3 gene, a gene for an embryonic transcription factor, is controlled by a retinoic acid repressible enhancer. EMBO J. 1991;10: 2997–3005. doi:10.1002/j.1460-2075.1991.tb07850.x

73. Agrawal P, Blinka S, Pulakanti K, Reimer MH, Stelloh C, Meyer AE, et al. Genome editing demonstrates that the-5 kb Nanog enhancer regulates Nanog expression by modulating RNAPII initiation and/or recruitment. J Biol Chem. 2021;296: 100189. doi:10.1074/jbc.RA120.015152

74. Rosa A, Brivanlou AH. A regulatory circuitry comprised of miR-302 and the transcription factors OCT4 and NR2F2 regulates human embryonic stem cell differentiation. EMBO J. 2011;30: 237–248. doi:10.1038/emboj.2010.319

75. Lu Y, Ren X, Wang Y, Han J. Post-translational modifications and secretion of Wnt proteins. Biomedical Journal of Scientific & Technical Research. 2018;9.

76. Gallet A, Rodriguez R, Ruel L, Therond PP. Cholesterol modification of hedgehog is required for trafficking and movement, revealing an asymmetric cellular response to hedgehog. Dev Cell. 2003;4: 191–204. doi:10.1016/s1534-5807(03)00031-5

77. Doubravska L, Krausova M, Gradl D, Vojtechova M, Tumova L, Lukas J, et al. Fatty acid modification of Wnt1 and Wnt3a at serine is prerequisite for lipidation at cysteine and is essential for Wnt signalling. Cellular Signalling. 2011;23: 837–848. doi:10.1016/j.cellsig.2011.01.007

78. Panáková D, Sprong H, Marois E, Thiele C, Eaton S. Lipoprotein particles are required for Hedgehog and Wingless signalling. Nature. 2005;435: 58–65. doi:10.1038/nature03504

79. Kaiser K, Gyllborg D, Procházka J, Salašová A, Kompaníková P, Molina FL, et al. WNT5A is transported via lipoprotein particles in the cerebrospinal fluid to regulate hindbrain morphogenesis. Nat Commun. 2019;10: 1498. doi:10.1038/s41467-019-09298-4

80. Farese RV, Ruland SL, Flynn LM, Stokowski RP, Young SG. Knockout of the mouse apolipoprotein B gene results in embryonic lethality in homozygotes and protection against diet-induced hypercholesterolemia in heterozygotes. Proceedings of the National Academy of Sciences. 1995;92: 1774–1778. doi:10.1073/pnas.92.5.1774

81. Tanaka Y, Okada Y, Hirokawa N. FGF-induced vesicular release of Sonic hedgehog and retinoic acid in leftward nodal flow is critical for left–right determination. Nature. 2005;435: 172–177. doi:10.1038/nature03494

82. Zhang XM, Ramalho-Santos M, McMahon AP. Smoothened mutants reveal redundant roles for Shh and Ihh signaling including regulation of L/R asymmetry by the mouse node. Cell. 2001;105: 781–792.

83. Okada Y, Takeda S, Tanaka Y, Belmonte J-CI, Hirokawa N. Mechanism of Nodal Flow: A Conserved Symmetry Breaking Event in Left-Right Axis Determination. Cell. 2005;121: 633– 644. doi:10.1016/j.cell.2005.04.008

84. Li Z, Schulz MH, Look T, Begemann M, Zenke M, Costa IG. Identification of transcription factor binding sites using ATAC-seq. Genome Biology. 2019;20: 45. doi:10.1186/s13059-019-1642-2

85. Avilion AA, Nicolis SK, Pevny LH, Perez L, Vivian N, Lovell-Badge R. Multipotent cell lineages in early mouse development depend on SOX2 function. Genes Dev. 2003;17: 126–140. doi:10.1101/gad.224503

86. DeVeale B, Brokhman I, Mohseni P, Babak T, Yoon C, Lin A, et al. Oct4 Is Required ∼E7.5 for Proliferation in the Primitive Streak. PLOS Genetics. 2013;9: e1003957. doi:10.1371/journal.pgen.1003957

87. Aires R, Jurberg AD, Leal F, Nóvoa A, Cohn MJ, Mallo M. Oct4 Is a Key Regulator of Vertebrate Trunk Length Diversity. Dev Cell. 2016;38: 262–274. doi:10.1016/j.devcel.2016.06.021

88. Graham V, Khudyakov J, Ellis P, Pevny L. SOX2 Functions to Maintain Neural Progenitor Identity. Neuron. 2003;39: 749–765. doi:10.1016/S0896-6273(03)00497-5

89. Amin S, Neijts R, Simmini S, van Rooijen C, Tan SC, Kester L, et al. Cdx and T Brachyury Co-activate Growth Signaling in the Embryonic Axial Progenitor Niche. Cell Reports. 2016;17: 3165–3177. doi:10.1016/j.celrep.2016.11.069

90. Bhattacharya A, Deng JM, Zhang Z, Behringer R, de Crombrugghe B, Maity SN. The B subunit of the CCAAT box binding transcription factor complex (CBF/NF-Y) is essential for early mouse development and cell proliferation. Cancer Res. 2003;63: 8167–8172.

91. Lu F, Liu Y, Inoue A, Suzuki T, Zhao K, Zhang Y. Establishing chromatin regulatory landscape during mouse preimplantation development. Cell. 2016;165: 1375–1388. doi:10.1016/j.cell.2016.05.050

92. Pollard KS, Hubisz MJ, Rosenbloom KR, Siepel A. Detection of nonneutral substitution rates on mammalian phylogenies. Genome Res. 2010;20: 110–121. doi:10.1101/gr.097857.109

93. Kurokawa R, Söderström M, Hörlein A, Halachmi S, Brown M, Rosenfeld MG, et al. Polarity-specific activities of retinoic acid receptors determined by a co-repressor. Nature. 1995;377: 451–454. doi:10.1038/377451a0

94. Berenguer M, Meyer KF, Yin J, Duester G. Discovery of genes required for body axis and limb formation by global identification of retinoic acid–regulated epigenetic marks. Su Y-H, editor. PLoS Biol. 2020;18: e3000719. doi:10.1371/journal.pbio.3000719

95. Yoon JK, Wold B. The bHLH regulator pMesogenin1 is required for maturation and segmentation of paraxial mesoderm. Genes Dev. 2000;14: 3204–3214. doi:10.1101/gad.850000

96. Chalamalasetty RB, Garriock RJ, Dunty WC, Kennedy MW, Jailwala P, Si H, et al. Mesogenin 1 is a master regulator of paraxial presomitic mesoderm differentiation. Development. 2014;141: 4285–4297. doi:10.1242/dev.110908

97. Waymack R, Fletcher A, Enciso G, Wunderlich Z. Shadow enhancers can suppress input transcription factor noise through distinct regulatory logic. Wittkopp PJ, Crocker J, editors. eLife. 2020;9: e59351. doi:10.7554/eLife.59351

98. Perry MW, Boettiger AN, Levine M. Multiple enhancers ensure precision of gap gene-expression patterns in the Drosophila embryo. Proc Natl Acad Sci U S A. 2011;108: 13570– 13575. doi:10.1073/pnas.1109873108

99. Bothma JP, Garcia HG, Ng S, Perry MW, Gregor T, Levine M. Enhancer additivity and non-additivity are determined by enhancer strength in the Drosophila embryo. Krumlauf R, editor. eLife. 2015;4: e07956. doi:10.7554/eLife.07956

100. El-Sherif E, Levine M. Shadow Enhancers Mediate Dynamic Shifts of Gap Gene Expression in the Drosophila Embryo. Current Biology. 2016;26: 1164–1169. doi:10.1016/j.cub.2016.02.054

101. Osterwalder M, Barozzi I, Tissières V, Fukuda-Yuzawa Y, Mannion BJ, Afzal SY, et al. Enhancer redundancy provides phenotypic robustness in mammalian development. Nature. 2018;554: 239–243. doi:10.1038/nature25461

102. Cunningham TJ, Lancman JJ, Berenguer M, Dong PDS, Duester G. Genomic Knockout of Two Presumed Forelimb Tbx5 Enhancers Reveals They Are Nonessential for Limb Development. Cell Rep. 2018;23: 3146–3151. doi:10.1016/j.celrep.2018.05.052

103. Frankel N, Davis GK, Vargas D, Wang S, Payre F, Stern DL. Phenotypic robustness conferred by apparently redundant transcriptional enhancers. Nature. 2010;466: 490–493. doi:10.1038/nature09158

104. Perry MW, Boettiger AN, Bothma JP, Levine M. Shadow enhancers foster robustness of Drosophila gastrulation. Curr Biol. 2010;20: 1562–1567. doi:10.1016/j.cub.2010.07.043

105. Antosova B, Smolikova J, Klimova L, Lachova J, Bendova M, Kozmikova I, et al. The Gene Regulatory Network of Lens Induction Is Wired through Meis-Dependent Shadow Enhancers of Pax6. PLOS Genetics. 2016;12: e1006441. doi:10.1371/journal.pgen.1006441

106. Wang H, Yang H, Shivalila CS, Dawlaty MM, Cheng AW, Zhang F, et al. One-Step Generation of Mice Carrying Mutations in Multiple Genes by CRISPR/Cas-Mediated Genome Engineering. Cell. 2013;153: 910–918. doi:10.1016/j.cell.2013.04.025

107. Casaca A, Nóvoa A, Mallo M. Hoxb6 can interfere with somitogenesis in the posterior embryo through a mechanism independent of its rib-promoting activity. Development. 2016;143: 437–448. doi:10.1242/dev.133074

108. Hogan B, Beddington R, Constantini F. Manipulating the Mouse Embryo: A Laboratory Manual. 4th ed. Cold Spring Harbor; 1994. Available: https://www.cshlpress.com/default.tpl?cart=1658310020248806022&fromlink=T&linkaction=full&linksortby=oop_title&--eqSKUdatarq=982

109. Tekko T, Lozovska A, Nóvoa A, Mallo M. Assessing Myf5 and Lbx1 contribution to carapace development by reproducing their turtle-specific signatures in mouse embryos. Dev Dyn. 2022. doi:10.1002/dvdy.502

110. Picelli S, Faridani OR, Björklund ÅK, Winberg G, Sagasser S, Sandberg R. Full-length RNA-seq from single cells using Smart-seq2. Nat Protoc. 2014;9: 171–181. doi:10.1038/nprot.2014.006

111. Dobin A, Davis CA, Schlesinger F, Drenkow J, Zaleski C, Jha S, et al. STAR: ultrafast universal RNA-seq aligner. Bioinformatics. 2013;29: 15–21. doi:10.1093/bioinformatics/bts635

112. Love MI, Huber W, Anders S. Moderated estimation of fold change and dispersion for RNA-seq data with DESeq2. Genome Biology. 2014;15: 550. doi:10.1186/s13059-014-0550-8

113. Ashburner M, Ball CA, Blake JA, Botstein D, Butler H, Cherry JM, et al. Gene ontology: tool for the unification of biology. The Gene Ontology Consortium. Nat Genet. 2000;25: 25–29. doi:10.1038/75556

114. Mi H, Muruganujan A, Ebert D, Huang X, Thomas PD. PANTHER version 14: more genomes, a new PANTHER GO-slim and improvements in enrichment analysis tools. Nucleic Acids Res. 2019;47: D419–D426. doi:10.1093/nar/gky1038

115. Correia RB, Barrat A, Rocha LM. The metric backbone preserves community structure and is a primary transmission subgraph in contact networks. bioRxiv; 2022. p. 2022.02.02.478784. doi:10.1101/2022.02.02.478784

116. Divate M, Cheung E. GUAVA: A Graphical User Interface for the Analysis and Visualization of ATAC-seq Data. Frontiers in Genetics. 2018;9. Available: https://www.frontiersin.org/articles/10.3389/fgene.2018.00250

117. Robinson JT, Thorvaldsdóttir H, Winckler W, Guttman M, Lander ES, Getz G, et al. Integrative genomics viewer. Nat Biotechnol. 2011;29: 24–26. doi:10.1038/nbt.1754

118. Kulakovskiy IV, Vorontsov IE, Yevshin IS, Sharipov RN, Fedorova AD, Rumynskiy EI, et al. HOCOMOCO: towards a complete collection of transcription factor binding models for human and mouse via large-scale ChIP-Seq analysis. Nucleic Acids Research. 2018;46: D252–D259. doi:10.1093/nar/gkx1106

119. Jurberg AD, Aires R, Varela-Lasheras I, Nóvoa A, Mallo M. Switching Axial Progenitors from Producing Trunk to Tail Tissues in Vertebrate Embryos. Developmental Cell. 2013;25: 451– 462. doi:10.1016/j.devcel.2013.05.009

120. Aires R, de Lemos L, Nóvoa A, Jurberg AD, Mascrez B, Duboule D, et al. Tail Bud Progenitor Activity Relies on a Network Comprising Gdf11, Lin28, and Hox13 Genes. Developmental Cell. 2019;48: 383–395.e8. doi:10.1016/j.devcel.2018.12.004

121. Lickert H, Kispert A, Kutsch S, Kemler R. Expression patterns of Wnt genes in mouse gut development. Mechanisms of Development. 2001;105: 181–184. doi:10.1016/S0925-4773(01)00390-2

122. Vinagre T, Moncaut N, Carapuço M, Nóvoa A, Bom J, Mallo M. Evidence for a myotomal Hox/Myf cascade governing nonautonomous control of rib specification within global vertebral domains. Dev Cell. 2010;18: 655–661. doi:10.1016/j.devcel.2010.02.011

123. Wang Y, Song F, Zhang B, Zhang L, Xu J, Kuang D, et al. The 3D Genome Browser: a web-based browser for visualizing 3D genome organization and long-range chromatin interactions. Genome Biology. 2018;19: 151. doi:10.1186/s13059-018-1519-9

